# Pre-injury subchronic stress confers sex-specific protection against pain-associated symptoms in osteoarthritic mice

**DOI:** 10.64898/2026.02.27.708540

**Authors:** Roxana Florea, Samuel Singleton, Laura Andreoli, Sara Hestehave, Christopher J. Black, Sandrine M. Géranton

## Abstract

Exposure to stress in adulthood alters the manifestation of persistent pain, yet its role in chronic pain vulnerability remains unclear. Here, we report that sub-chronic restraint stress experienced two weeks before unilateral osteoarthritis (OA) induction *via* intra-articular mono-iodoacetate (MIA) promoted the emergence of a resilient-like phenotype in adult male mice. This was evidenced by decreased MIA-induced mechanical hypersensitivity, improved gait dynamics and lower anxiety-like behaviour. In contrast, stress exposed females exhibited augmented pain-associated symptoms. In males, sub-chronic stress mitigated several MIA-induced molecular changes, including reduced adult hippocampal neurogenesis, increased glucocorticoid receptor levels in the hypothalamic paraventricular nucleus (PVN) and elevated c-Fos expression in deep spinal laminae, periaqueductal gray (PAG) and PVN. RNA sequencing suggested that restraint stress in males primed the spinal cord for an exacerbated GABAergic response post-MIA and pre-empted some pro-nociceptive MIA-induced transcriptional changes. However, the combination of stress and injury disrupted longevity-associated programs and shifted neurons toward a stress-vulnerable state. These findings reveal that prior stress exposure can modulate the long-term pain experience in osteoarthritis, with sexually dimorphic outcomes. Importantly, these results challenge the notion of stress as inherently maladaptive and underscore its potential to foster resilience to chronic pain.

## 1. Introduction

When faced with internal or external challenges – collectively referred to as stressors – organisms mount an evolutionarily conserved stress response that rapidly enhances alertness, arousal and cognitive processing to promote survival. Acute, but intense stress can reduce pain perception in both clinical and preclinical settings through a phenomenon known as stress-induced analgesia^1–5^. Whilst this short-lasting enhancement to endogenous analgesic tone enables the organisms to maintain functionality and escape danger even when injured, when prolonged, the stress response can increase the risk of developing a wide range of affective and somatic disorders, including chronic pain^6–8^. In preclinical settings, prolonged exposure to stress increases somatic sensitivity to chemical, mechanical, and thermal stimuli, through a phenomenon known as stress-induced hyperalgesia^9^. Although heightened sensitivity could serve an adaptive function in certain contexts^10^, it resembles the maladaptive plasticity of nociceptive circuits that underlies many pathological pain conditions. Therefore, understanding the neurobiological mechanisms underlying stress-related plasticity in the central nervous system could provide unique insights into the processes that contribute to pathological pain.

As one of leading causes of chronic pain worldwide, osteoarthritis (OA) is an increasingly prevalent degenerative disease of the joints for which effective therapies are still lacking^11^. Although symptoms such as joint stiffness, swelling, and reduced function are characteristic of OA and can impair quality of life, it is pain that most commonly drives patients to seek medical care. However, effective pain management options remain limited for these patients, partly because the underlying mechanisms that generate and maintain pain remain poorly understood^12–14^.

Clinical observations indicate that OA evolves into a complex, multidimensional pain condition wherein structural joint pathology alone cannot account for the variability in pain reports. This growing recognition highlights the urgent need to integrate psychological, cognitive, and socioeconomic factors to better understand pain heterogeneity in OA^15–17^. Indeed, mood disorders such as anxiety and depression are prevalent among individuals with OA^18^, while cognitive factors such as catastrophizing, low self-efficacy, negative coping, and fear of movement are associated with pain^19–22^. While these findings highlight the strong interplay between pain and affective states, it remains unclear how pre-existing individual characteristics – including those shaped by prior stress exposure – may influence the trajectory and variability of pain experienced throughout OA progression.

To address this gap, we investigated the effects of subchronic restraint stress on the subsequent pain experience in the mono-iodoacetate (MIA) mouse model of OA. Much of the existing research has focused on prolonged and/or severe stress paradigms, rather than milder and predictable stressors such as those encountered in daily life, leaving a gap in the stress literature. Accordingly, in our study, male and female mice were exposed to three consecutive days of 1h restraint stress (RS). We and others have characterized the RS paradigm and shown that it induces short-lived mechanical hypersensitivity – no longer than 10 days - but long-lasting epigenetic changes ^23^. The stress exposure was followed by an intra-articular MIA injection two weeks later. Our findings reveal that prior stress conferred resilience to joint pain in male mice, while it exacerbated the pain- and promoted the emergence of anxiety-like behaviours in females. The resilient-like phenotype observed in male mice was accompanied by altered spinal GABAergic signalling and reduced neuronal activity in CNS regions implicated in stress and pain processing (***fig. S1, graphical abstract***).

## 2. Results

All statistical tests and extended analysis results are presented in **Supplementary Table S1.**

### 2.1 Subchronic stress exposure reduces mechanical hypersensitivity and functional deficit induced by unilateral knee arthritis in male, but not female mice

Two weeks after the first RS exposure, mice received a unilateral knee injection of MIA (day 0) (***Fig. 1A***). In males, both stressed and non-stressed mice displayed a significant bilateral reduction in paw withdrawal thresholds (PWTs) compared to control mice that lasted at least 27 days (Ipsi: ***Fig. 1B*:** RM ANOVA D-1 to 27: ‘Treatment’ F_2,19_=8.18, *P* = 0.0027; Šídák post-hoc: Control *vs* MIA: *P* = 0.003; Control *vs* RS+MIA: *P* = 0.018; ***Fig. 1C***: ANOVA: F_2,19_=9.11, *P* = 0.0017; Contra: ***Fig. 1D***: RM ANOVA D-1 to 27: ‘Treatment’ F_2,19_=11.67, *P* = 0.005; Šídák post-hoc: Control *vs* MIA: *P* < 0.001; Control *vs* RS+MIA: *P* = 0.052). The increase in cutaneous sensitivity was delayed on the ipsilateral side in mice exposed to RS compared to non-stressed mice (***Fig. 1B***: RM ANOVA D-1 to 8: ‘Treatment’ F_2,19_=5.49, *P* = 0.013; Šídák post-hoc: Control *vs* MIA: *P* = 0.011; Control *vs* RS+MIA: *P* = 0.303), while on the contralateral side RS had a more prominent effect, substantially reducing the MIA-induced mechanical hypersensitivity (***Fig. 1D*:** RM ANOVA: D4 to 23: ‘Treatment’ F_2,19_=14.90, *P* < 0.001; Šídák post-hoc: MIA *vs* RS+MIA: *P* = 0.039; ***Fig. 1E***: ANOVA: D-1 to 27, F_2,19_=14.32, *P* = 0.0002).

**Figure 1.**
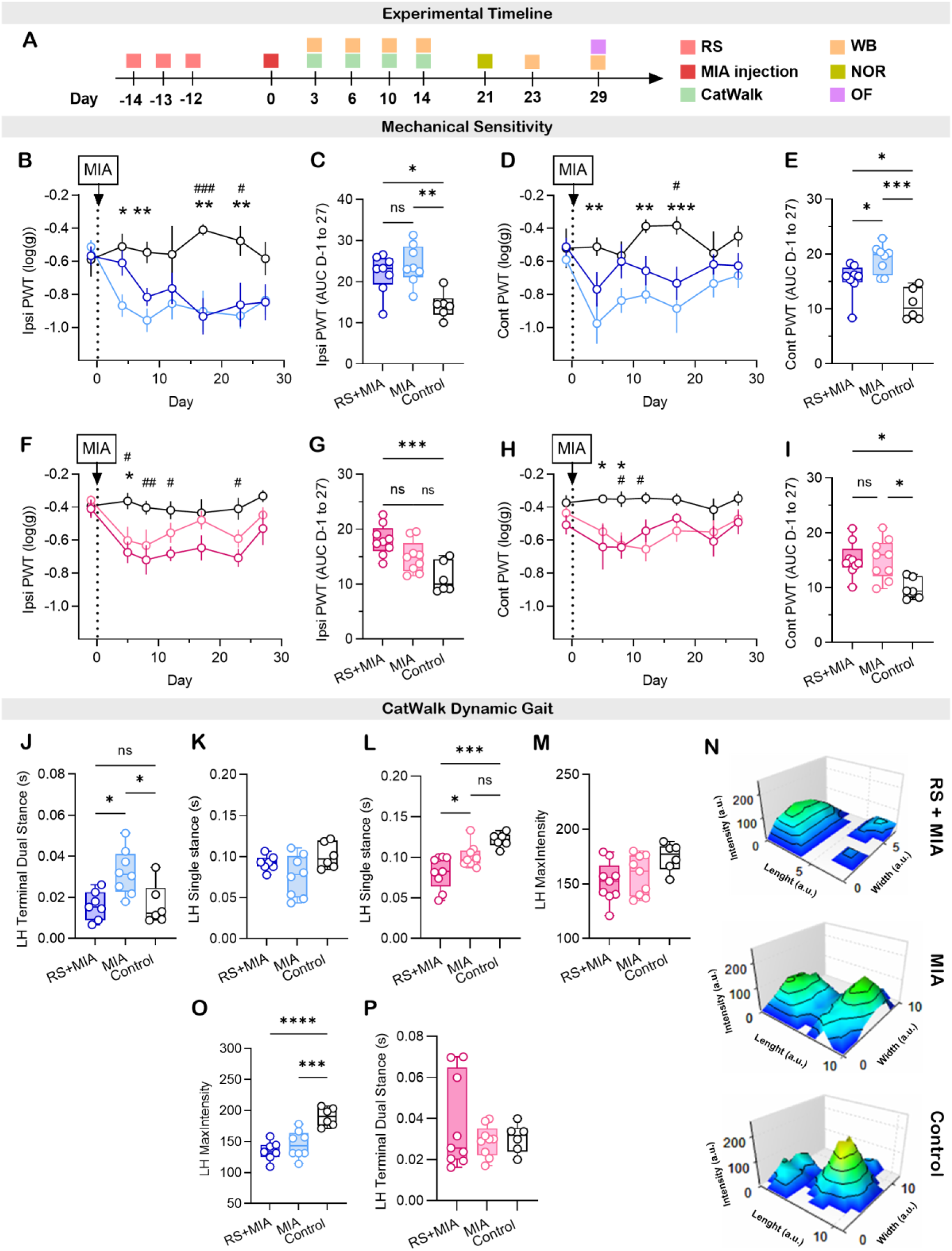
Subchronic restraint stress attenuates MIA-induced mechanical allodynia and dynamic gait impairments at peak deficit in male, but not female mice. (A) Experimental timeline. RS – restraint stress; MIA – monoiodoacetate; WB – weight bearing; NOR – novel object recognition test; OF – open field test. (B, C) Time course (B) and AUC (C, day -1 to 27) of mechanical sensitivity at the ipsilateral hindpaw in male mice. (D, E) Time course (D) and AUC (E, day -1 to 27) of mechanical sensitivity at the contralateral hindpaw in male mice. *N* = 8/8/6. \**P*<0.05, \*\**P*<0.01 MIA *vs* Control; ^#^*P*<0.05, ^###^*P*<0.001 RS+MIA *vs* Control. (F, G) Time course (F) and AUC (G, day -1 to 27) of mechanical sensitivity at the ipsilateral hindpaw in female mice. (H, I) Time course (H) and AUC (I, day -1 to 27) of mechanical sensitivity at the contralateral hindpaw in female mice. *N* = 9/9/6. \**P*<0.05 MIA *vs* Control; ^#^*P*<0.05, ^##^*P*<0.01 RS+MIA *vs* Control. (J-M, O, P) Individual bar plots of dynamic gait parameters at day six post MIA injection in male. LH: left hind paw = ipsilateral hindpaw. (J, K, O; *N* = 7/8/6) and female (L, M, P; *N* = 9/9/6) mice. (N) 3D footprint intensities of the ipsilateral hindpaw of typical female mice from each treatment group (note that in control mice, the largest peak corresponds to heel pressure with smaller peaks reflecting digit contact. In MIA mice, heel intensity is reduced and digit peaks partially merge, while in RS+MIA mice heel pressure is markedly diminished and digit peaks collapse into a single, smaller area, reflecting reduced paw contact and weight bearing on the injured hindlimb; scale: 0-255, a.u. – arbitrary unit). B, D, F, H data expressed as mean ± SEM.

MIA also induced bilateral mechanical hypersensitivity in non-stressed female mice (***Fig.1* *F-I; Table S1***). However, in contrast to the findings in males, stress exposure exacerbated the MIA-induced mechanical allodynia at the ipsilateral hindpaw in females (***Fig. 1F*:** RM ANOVA D-1 to 27: ‘Treatment’ F_2,21_=8.95, *P* = 0.0015; Šídák post-hoc: Control *vs* MIA: *P* = 0.079; Control *vs* RS+MIA: *P* = 0.001; MIA *vs* RS+MIA: *ns; LSD* post-hoc: Control *vs* MIA: *P* = 0.027; Control *vs* RS+MIA: *P* < 0.001; MIA *vs* RS+MIA: P=0.05; ***Fig. 1G***: ANOVA: F_2,21_=10.36, *P* = 0.0007), with no obvious effect on the contralateral PWTs (***Fig. 1H***, RM ANOVA D-1 to 27: ‘Treatment’ F_2,21_=6.83, *P* = 0.005; Šídák post-hoc: RS+MIA *vs* Control: *P* = 0.008; MIA *vs* Control: *P* = 0.012; MIA *vs* RS+MIA: *ns*; ***Fig. 1I***: ANOVA D-1 to D27: F_2,21_=6.51, *P* = 0.006).

A comprehensive analysis of gait behaviour and static weight bearing was conducted on the same animals using the automated CatWalk system and incapacitance assay, respectively. Overall, the MIA injection led to transient changes in dynamic gait behaviours that peaked at day six after injury for both male and female mice (***fig. S2A-D***). Exposure to RS had a significant but complex and sex-specific impact on dynamic gait parameters. Evaluation at peak deficit (day six) revealed that stress exposure reversed the MIA-induced impact on selected parameters in male mice. Specifically, RS reversed the MIA-induced alterations in ground contact time of the injured hindleg which suggested a reluctance of MIA treated mice to promptly move the affected hindleg during locomotion (terminal dual stance, ***Fig. 1J***, F_2,18_=7.06, *P* = 0.0055); with a similar trend observed for the contact duration of the injured hindpaw (single stance time, ***Fig. 1K***, F_2,18_=3.22, *P* = 0.064) and for the proportion of time during the step cycle during which opposite fore- and hindlimbs are simultaneously in contact with the ground (diagonal support; ***fig S2E)***. In contrast, in female mice, prior stress exposure significantly decreased the contact duration of the injured hindpaw with the runway compared to both control and MIA mice (***Fig. 1L***, F_2,21_=11.44, *P* = 0.0004), with a similar trend found for the maximum pressure (maximum intensity, ***Fig. 1M***, F_2,21_=3.248, *P* = 0.059, ***N***). These observations could not be attributed to potential differences in exploratory activity, as the running speeds along the corridor were similar across all groups in male (***fig. S2F***) and female mice (***fig. S2G***). Although control males and females may have differed for several parameters due to differences in running speed and morphometric characteristics between sexes, sexes were not directly compared and speed did not differ within sex subgroups. Therefore, this did not confound our findings on the impact of stress exposure on dynamic gait in males and females. Finally, not all changes in gait parameters were modulated by RS at day six after MIA (***Fig. 1O***, F_2,19_=18.49, *P* < 0.0001, ***P***). Similarly, RS did not significantly impact the MIA-induced static weight bearing asymmetry (***fig. S2H***, ***I***).

Together, these findings suggest that exposure to subchronic restraint stress attenuated the MIA-induced mechanical hypersensitivity and gait deficit in male mice, but had an opposite effect in female mice.

### 2.2 Subchronic stress exposure prevents the development of MIA-induced anxiety-like behaviour in male, but not female mice

Anxiety-like behaviour was assessed using the open field test at one month (day 29) after the MIA injection. In male mice, MIA promoted the emergence of anxiety-like behaviours, as evidenced by a significant reduction in both the probability of time spent and distance travelled in the center of the arena (***Fig. 2A***, F_2,18_=5.15, *P* = 0.017, ***B***: F_2,18_=5.97, *P* = 0.0103), outcomes reversed by prior RS exposure. There were no obvious differences observed in activity in males within the peripheral zones (***Fig. 2C-E***, ***fig. S3A***). In female mice however, prior RS exposure promoted the emergence of anxiety-like behaviours as indicated by an increased probability of RS+MIA, but not MIA, mice to occupy the ‘Far Away from Center’ zone compared to controls (***Figure 2F-I, H***: F_2,20_=3.8, *P* = 0.0399; ***I***: F_2,20_=4.36, *P* = 0.0269). No significant differences were found for the total distance travelled in the open field arena by female mice (***Fig. 2G***, ***fig. S3B***) and all observations were confirmed using time spent in each of the OF zones as outcome measures (***fig. S4A-F***).

**Figure 2.**
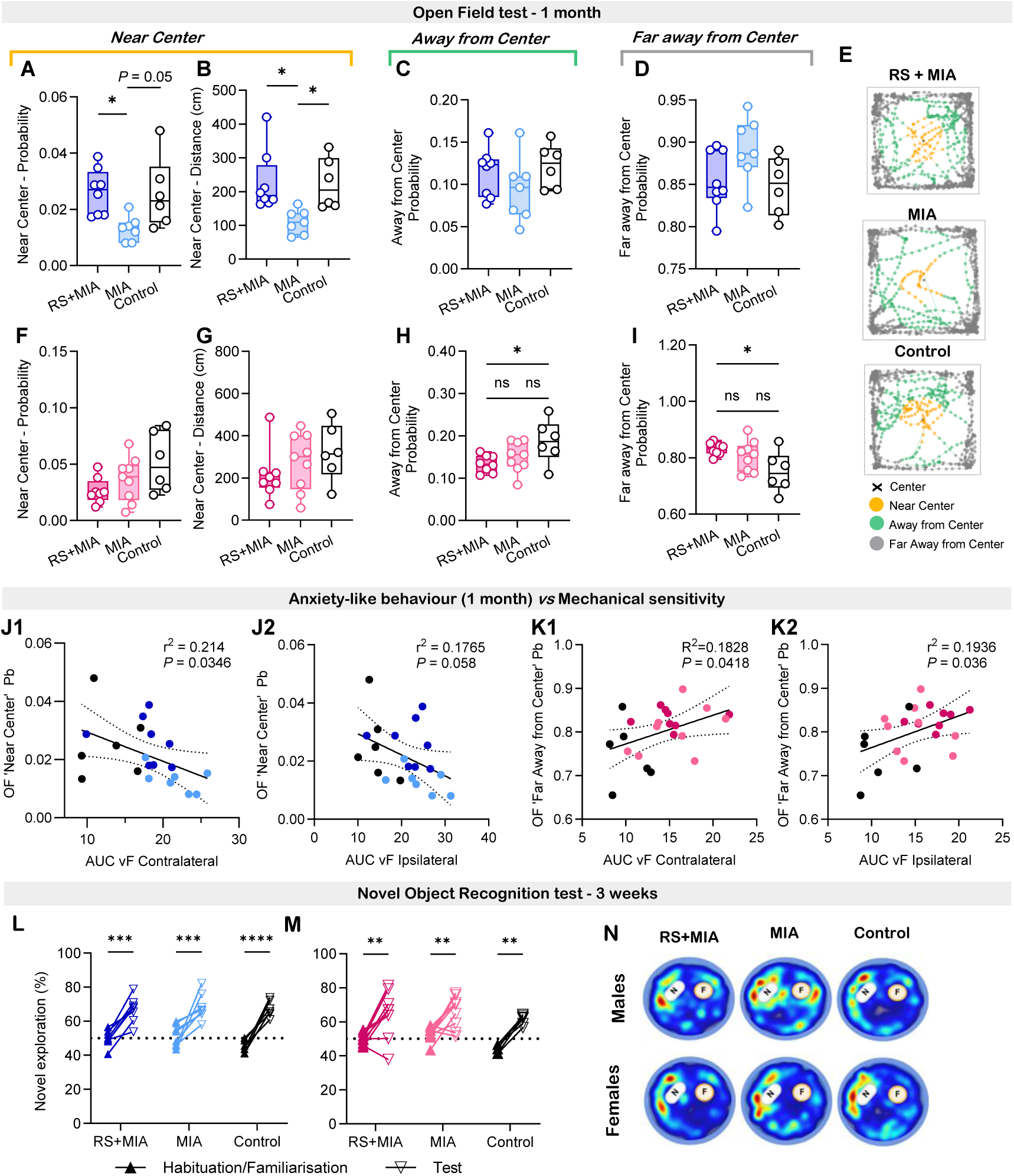
Subchronic stress exposure reverses MIA-induced anxiety-like behaviour in male mice while it promotes its development in female mice. (**A-D**) Quantification of anxiety-like behaviour of male mice in the ‘Near Center’ (**A**, **B**), ‘Away from Center’ (**C**), and ‘Far away from Center’ (**D**) zones of the open field arena. *N* = 8/7/6. (**E**) Representative scatter plots illustrating the spatial distribution of male mice’s movement across the defined zones of the open field arena. (**F**-**I**) Quantification of anxiety-like behaviour of female mice in the ‘Near Center’ (**F**, **G**), ‘Away from Center’ (**H**), and ‘Far away from Center’ (**I**) zones of the arena. *N* = 8/9/6. (**J1**) Correlation analysis between the probability of mice being located in the ‘Near Center’ zone of the open field arena and the area under curve (AUC; D-1 to 27 after MIA) for mechanical allodynia at the contralateral hindpaw for male mice. *N* = 21. (**J2**) Correlation analysis between hindpaw mechanical sensitivity and the probability of male mice being in the ‘Near Center’ zone of the open field arena and the area under curve (AUC; D-1 to 27 after MIA) for mechanical allodynia at the ipsilateral hindpaw for male mice. *N* = 21. (**K1**) Correlation analysis between hindpaw mechanical sensitivity and the probability of female mice being in the ‘Far Away from Center’ zone of the open field arena and the area under curve (AUC; D-1 to 27 after MIA) for mechanical allodynia at the contralateral hindpaw for male mice. *N* = 23. (**K2**) Correlation analysis between the probability of mice being located in the ‘Far Away from Center’ zone of the open field arena and the area under curve (AUC; D-1 to 27 after MIA) for mechanical allodynia at the ipsilateral hindpaw for female mice. *N* = 23. (**L**-**N**) Time spent exploring the novel object expressed as percentage of the total amount of time spent exploring the familiar and novel objects in male (**L**, *N* = 8/8/6) and female (**M**, *N* = 9/9/6) mice, and representative heatmaps (**N**, N – novel object; F – familiar object). \**P*<0.05, \*\**P*<0.01, \*\*\**P*<0.001, \*\*\*\**P*<0.0001. Solid lines in **J** and **K** indicate linear regression fits, while dashed lines the 95% CI.

Given the strong association between pain and mood disorders seen in patients with OA, we next conducted a correlation analysis between pain- and anxiety-like behaviours in male and female mice. Focusing on the behavioural parameters most affected by RS in each sex, we found a significant correlation in males between the AUC of contralateral PWTs and the probability of spending time the center of the open field arena (***Fig. 2J1***, F_1,19_=5.18, *P* = 0.0346), with a similar trend observed for ipsilateral hindpaw responses (***Fig. 2J2***). In females, significant correlations were identified between the probability of spending time in the border zone of the arena and both ipsilateral (***Fig. 2K1***, F_1,21_=4.7, *P* = 0.0418) and contralateral hindpaw sensitivity to mechanical stimulation (***Fig. 2K2,*** F_1,21_=5.04, *P* = 0.0357).

We also explored the impact of prior stress exposure on cognitive functioning and performed an assessment of novelty preference using the novel object recognition assay at three weeks after the MIA injection. We found that both male and female cohorts spent significantly more time exploring the novel object compared to the familiar one, irrespective of the treatment group, suggesting no impact of MIA or RS exposure on cognitive function at 3 weeks post MIA (***Fig. 2L-N*,** RM ANOVA: **L**: ‘Time’ F_1,19_=90.9, *P* < 0.0001; **M**: ‘Time’ F_1,21_=44.52, *P* < 0.0001).

Lastly, quantification of body weight at the time of anaesthesia revealed that the MIA injection attenuated body weight gain in male mice which was no mitigated by RS exposure (***fig. S3C***). No group differences were observed in the female cohort (***fig. S3D***).

Collectively, these findings suggest that subchronic restraint stress mitigated the development of anxiety-like behaviours after the knee MIA injection in male mice, with anxiety-like behaviour correlating with mechanical hypersensitivity away from the injury. In female mice however, RS promoted the emergence of anxiety-like behaviours which also correlated with the secondary hyperalgesia.

### 2.3 At one month after the initiation of the arthritic state, prior subchronic stress exposure prevents MIA-induced changes in hippocampal neurogenesis and GR expression in the PVN of male mice

To investigate the molecular mechanisms underlying the observed behavioural outcomes, brain and spinal cord tissues were collected from the male and female mice that had been behaviourally characterised (***Fig.1***, ***2***) at one month after the MIA injection. As both chronic pain and stress can disrupt hippocampal plasticity^23,24^, we examined doublecortin (DCX) expression to assess potential alterations in neurogenic processes under these conditions. Analysis of immunofluorescence staining in the dentate gyrus of the hippocampus showed a reduction in DCX expression following MIA injection in male mice, reduction that was prevented by prior RS exposure (***Fig. 3A***, F_2,17_=3.54, *P* = 0.050, ***B***). Supporting these observations, we found that RS alone increased DCX expression in the hippocampus of un-injured male mice (***fig S5A***). In contrast, no significant changes in DCX expression were detected in female mice at one month after the MIA injection (***Fig. 3C***).

**Figure 3.**
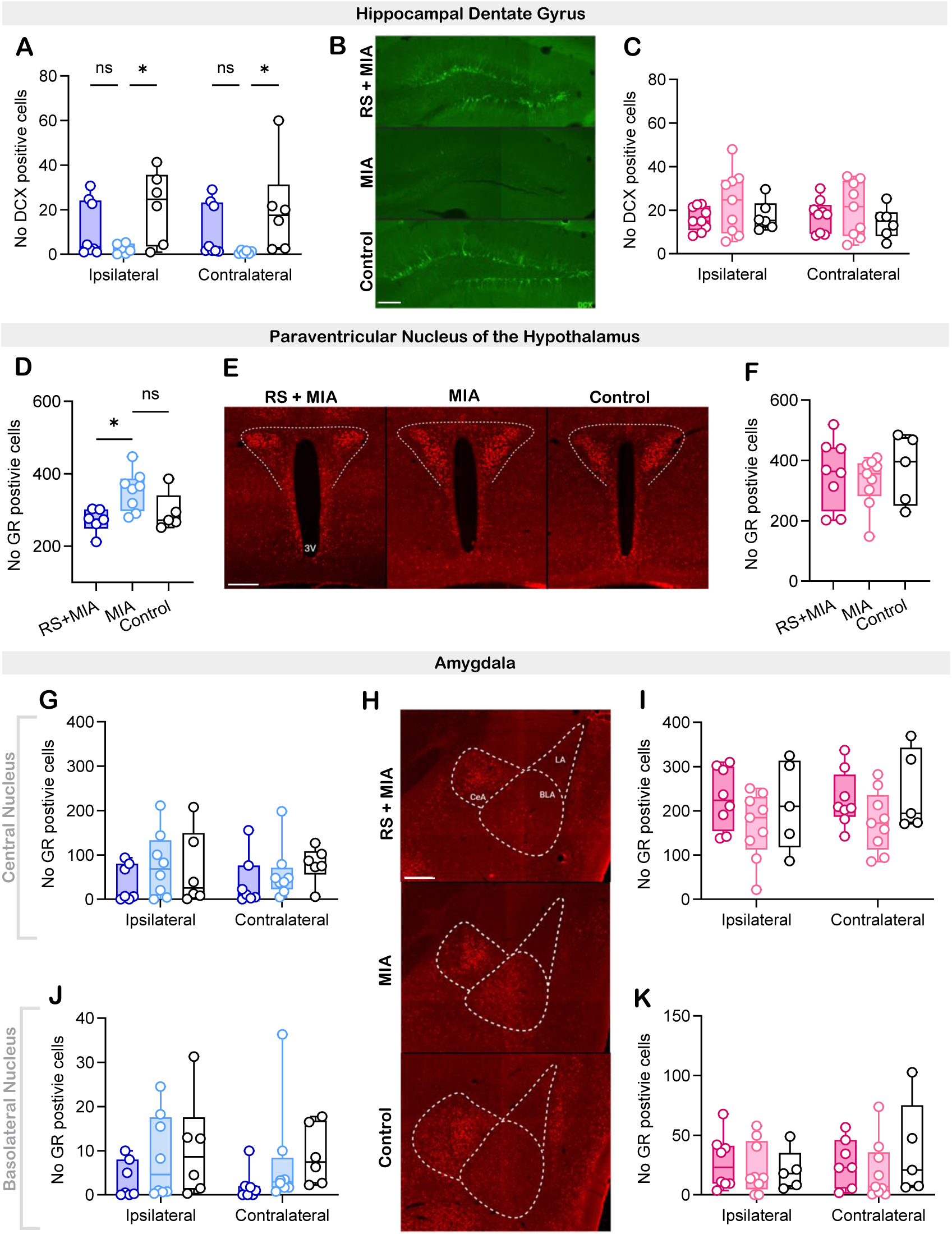
Subchronic restraint stress mitigates the MIA-induced changes in hippocampal neurogenesis and GR expression in the PVN in male mice at one month after injection. (**A, B**) Quantification of hippocampal neurogenesis (**A**, *N* = 8/6/6) and representative images of doublecortin (DCX) staining within the dentate gyrus of male mice (**B**, scale bar = 100μm). (**C**) Quantification of hippocampal neurogenesis in female mice (*N* = 9/9/6) (**D, E**) Quantification of glucocorticoid receptor (GR) expression in the paraventricular nucleus (PVN) of the hypothalamus (**D**, *N* = 6/8/5) and representative images of GR immunofluorescence within the PVN of male mice (**E**, scale bar = 200μm). (**F**) Quantification of GR positive cells in the hypothalamic PVN of female mice (*N* = 8/9/5). (**G, H**) Quantification of GR expression within the central nucleus (CeA) of amygdala in male mice (**G**, *N* = 7/8/6) and representative images (**H**; LA – lateral nucleus; scale bar = 200μm). Quantification of GR expression within CeA in females (**I**, *N* = 8/9/5). (**J**, **K**) Quantification of GR expression in the basolateral nucleus (BLA) of male (**J**, *N* = 7/8/6) and female (**K**, *N* = 8/9/5) mice.

Glucocorticoid receptor (GR) expression in limbic brain regions plays a critical role in modulating the HPA axis activity and shaping cognitive and affective responses to stress^25,26^. The PVN of the hypothalamus is a central hub that integrates external and internal stress signals, initiating HPA axis activation. In the PVN of male mice, GR expression in RS+MIA mice was lower than in MIA mice and not different from control mice, suggesting that prior stress restores GR expression toward a control-like state (***Fig. 3D***, F_2,16_=5.21, *P* = 0.0181; ***E***). In females however, no significant differences were observed (***Fig. 3F***). Additionally, no significant differences were observed for the number of GR positive cells in the central nucleus of the amygdala (CeA) in male (***Fig. 3G***, ***H***) or female (***Fig. 3I***) mice, nor in the basolateral nucleus (BLA, ***Fig. 3J***, ***K***, respectively). Similarly, quantification of GR fluorescence intensity in the hippocampus, a key negative regulator of HPA axis activity, revealed no significant group differences (***fig. S5B-D***).

Taken together, these findings indicate that RS, MIA and their combination elicited more prominent molecular alterations in stress-related central markers in male mice than in females. Given the novelty of the resilient-like phenotype observed in males, the subsequent molecular investigations were performed in males only.

### 2.4 Subchronic stress exposure reduces MIA-induced c-Fos expression in central pain and stress-regulating circuits and attenuates spontaneous nocifensive behaviours in male mice

We next sought to determine whether prior subchronic stress exposure in early adulthood had an impact on nociceptive circuits. To test this, spinal and supraspinal tissues were collected 2h after the knee injection of MIA for quantification of the immediate early gene product c-Fos^27^.

At the spinal cord level, a significant ‘Treatment’ effect was found for the number of c-Fos positive cells in the dorsal horn (LI-V) across laminae and spinal cord sides (***Fig. 4A***, 3-way ANOVA, factor ‘Treatment’ F_2,44_ = 6.87, *P* = 0.003, interactions: ‘Treatment x Laminae’: F_2,44_=8.82, *P* < 0.001). Post-hoc Šídák analysis revealed a small but significant increase in the number of c-Fos immunoreactive cells in superficial laminae I-II in MIA and RS+MIA mice compared with control mice (2-way ANOVA: ‘Treatment’ F_2,22_=5.12, *P*=0.015; Šídák post-hoc: MIA *vs* Control, *P* = 0.026; RS+MIA *vs* Control, *P* = 0.035), but there was no difference in c-Fos expression between MIA and RS+MIA mice. In contrast, in deeper laminae, the expression of c-Fos was significantly lower in RS+MIA *vs* MIA mice (2-way ANOVA: ‘Treatment’ F_2,22_=8.05, *P*=0.002; Šídák post-hoc: RS + MIA *vs* Control, *P* = 0.02; RS+MIA *vs* MIA, *P* = 0.004; complete stats report available in ***Table S1*** and further post-hoc analysis as plotted on ***Fig. 4A***).

**Figure 4.**
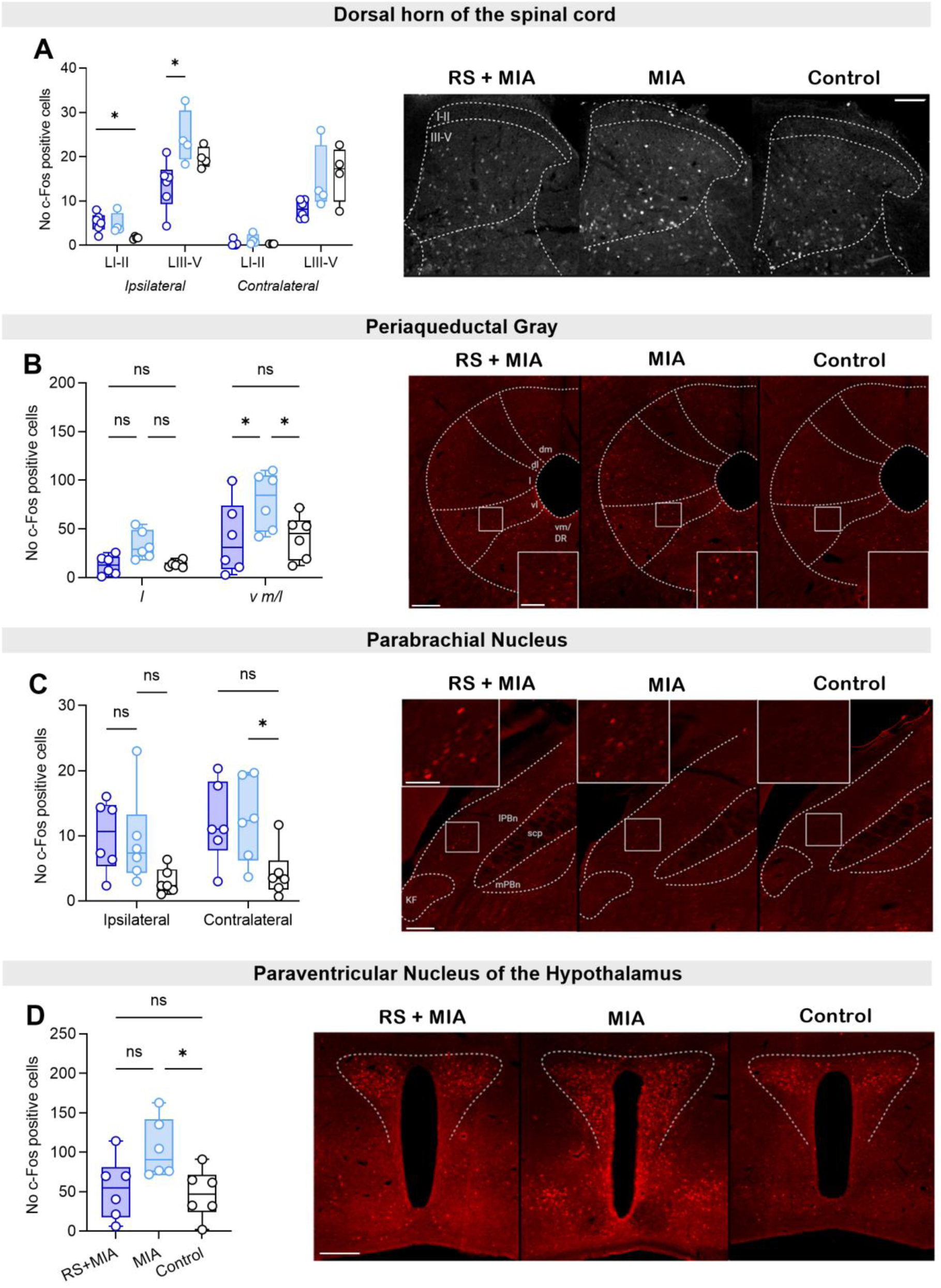
Restraint stress prevents MIA-induced c-Fos expression in male mice. (**A**) Quantification (left) and representative images (right; ipsilateral, scale bar = 100μm) of c-Fos immunostaining in the dorsal horn of the spinal cord. *N* = 6/4/4. (**B**) Quantification (left) and representative images (right; scale bar = 100μm; boxed areas scale bar = 50μm) of c-Fos immunostaining in the periaqueductal gray (PAG; dm – dorsomedial, dl – dorsolateral, l – lateral; vl – ventrolateral; v m/l – ventromedial/lateral). *N* = 6/6/6. (**C**) Quantification (left) and representative images (right; scale bar = 100μm; boxed areas scale bar = 50μm) of c-Fos immunostaining in the parabrachial nucleus (PBn; KF - Kölliker-Fuse nucleus, scp – superior cerebellar peduncle). *N* = 6/6/6. (**D**) Quantification (left) and representative images (right; scale bar = 200μm) of c-Fos immunostaining in the paraventricular nucleus of the hypothalamus. *N* = 6/6/6. \**P*<0.05.

In the brainstem, RS reversed the MIA-induced upregulation of c-Fos expression in the PAG (***Fig. 4B***, F_2,15_=4.6, *P* = 0.0276). A significant overall effect of MIA was found in the PBn, however post-hoc analysis comparing the individual groups failed to reveal a significant effect of prior RS exposure (***Fig. 4C***, F_2,15_=4.54, *P* = 0.0287). No significant differences in c-Fos expression between groups were observed in the RVM or the LC (***fig. S5E, F***). Finally, within the PVN of the hypothalamus, MIA mice, but not RS+MIA mice showed a significant increase in c-Fos expression compared to controls (***Fig. 4D***, F_2,15_=4.52, *P* = 0.0291).

We also sought to determine the impact of RS on the nociceptive responses induced by the intra-plantar injection of 5% formalin solution in male mice (***fig S6A***). Mice previously exposed to stress spent significantly less time attending the injured hindpaw in the second phase of the nocifensive behaviour compared to non-stressed mice (***fig. S6B***). At the molecular level, no changes in c-Fos expression were seen in any of the CNS regions investigated (***fig. S6C-F***).

Collectively, these findings demonstrate that RS mitigates the MIA-induced c-Fos expression in pain-and stress-regulating circuits at the early onset of joint pathology in male mice. Furthermore, the pain-relieving impact of RS is preserved in male mice exposed to a shorter-lasing algogenic compound.

### 2.5 The combination of RS and MIA elicits unique signalling pathways absent in MIA-only animals

Finally, we used RNA sequencing to establish whether transcriptional changes at spinal cord levels could explain the development of resilience induced by subchronic stress in male mice. For this, we compared gene expression across 4 groups of animals: a control group, a group only exposed to restraint stress, a group only exposed to MIA, and one group exposed to both RS and MIA (***Fig. 5A***). The spinal cord was divided into quadrants to explore separately MIA-induced ipsilateral from contralateral changes in the superficial dorsal horn.

**Figure 5.**
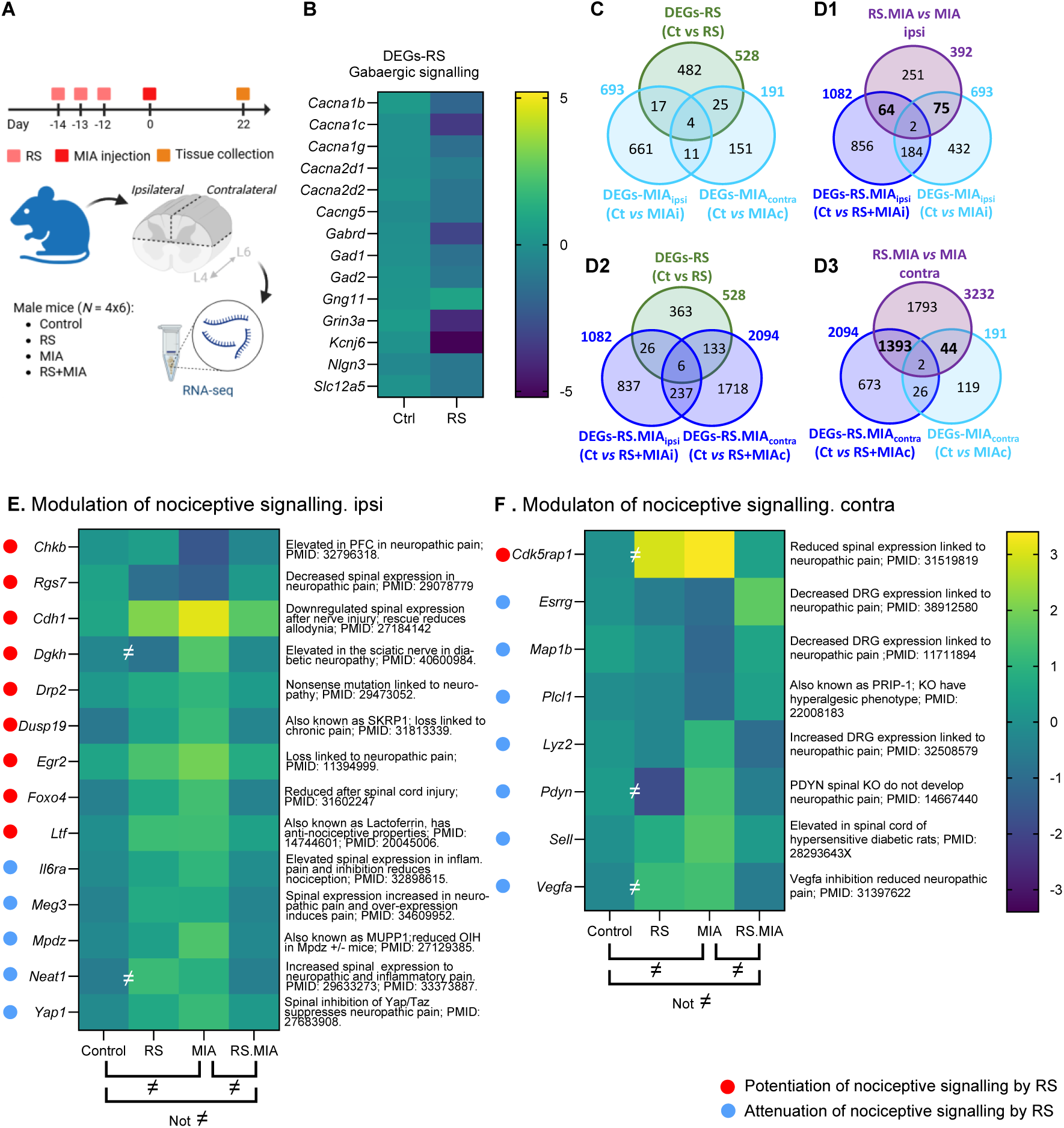
Exposure to RS reverses a number of MIA-induced transcriptional changes at spinal cord level. (**A**) Experimental timeline. (**B**) Heatmap showing differential expression of genes associated with GABAergic signalling following RS exposure. **(C, D,** Venn diagrams summarising DEGs between control (Ct), RS, MIA, and RS+MIA conditions illustrating additive or unique transcriptomic responses; (**E**, **F**) Heatmaps of nociceptive-related genes in the ipsilateral (**E**) and contralateral (**F**) spinal cord, showing distinct regulatory profiles. RS-associated potentiation or attenuation of nociceptive signalling is indicated with red and blue borders, respectively. “≠” denotes statistically significant differences across conditions.

First, we assessed the impact of RS exposure on spinal transcriptomes by comparing RS *vs* Control tissue using contralateral quadrants. We found 300 up-regulated and 228 down-regulated annotated genes (DEGs-RS). Pathway analysis using both Ingenuity Pathway Analysis (IPA) software and Enrichr revealed that the main regulated pathway was “GABAergic synapse/receptor signalling pathway” (IPA: -log(P)=2.68; Ratio+0.105; Z-score: 0.535; Enrichr: nominal *P* = 0.0024), with all but one gene downregulated following RS (***Fig. 5B, Table S2***; only upregulated gene: *Gng11*). Overall, these results suggested that the impact of sub-chronic stress exposure on the spinal transcriptome was long-lasting and lead to substantial changes in expression in genes associated with GABAergic signalling.

We next established the impact of MIA on gene expression in the ipsilateral or contralateral spinal cord through 2 separate contrasts: MIA ipsi *vs* Control ipsi (DEGs-MIA_ipsi_) and MIA contra *vs* Control contra (DEGs-MIA_contra_). There were 693 annotated DEGs on the ipsilateral side and 191 on the contralateral side. This suggested that MIA induced a greater number of transcriptional changes on the ipsilateral side than those induced by stress exposure, whereas fewer changes were observed on the contralateral side (***Fig. 5C***). On the ipsilateral side, we identified several pathways involved in osteoarthritis (including the osteoarthritis signalling pathway, PI3K/AKT, chondrocyte signalling, FAK signalling) confirming the validity of our osteoarthritis MIA model (***Table S3***). Fewer pathways were enriched on the contralateral side, but ‘osteoarthritis signalling’ stood out as significantly enriched (***Table S4***). 15 annotated DEGs were common to both contrasts and a total 46 DEGs regulated post-MIA across ipsi-and contra-lateral side were also regulated post-RS (***Fig. 5C***), with 4 DEGs common to all 3 contrasts (***fig. S7A***). These results suggested that, as expected, gene expression changes following unilateral knee joint injection of MIA differ between the ipsilateral and contralateral sides. Moreover, a subset of genes modulated by MIA may have been indirectly regulated *via* activation of the HPA axis, as seen with the overlap with RS-induced gene changes, with this effect occurring to a similar extent on both sides.

We then explored the impact of prior RS exposure to the MIA-induced changes in gene expression. 3176 annotated DEGs were identified in the contrasts RS.MIA *vs* Control (1082 DEGs-RS.MIA_ipsi_ and 2094 DEGs-RS.MIA_contra;_ ***Fig. 5D1****,**3***). As we were surprised by the extensive changes seen on the contralateral side compared with the ipsilateral side after combined exposure to RS and MIA, we ran further controls to assess sample integrity. However, visual inspection of multidimensional plots confirmed that RS.MIA samples clustered as expected, and the bulk RNA data revealed comparable proportions of each cell type between groups.

On the ipsilateral side (***Fig. 5D1***), 186 of the 1082 DEGs-RS.MIA_ipsi_ (17.2%) were also modulated by MIA in non RS-exposed animals: 184 showed the same level of modulation (*i.e.,* there was no differential expression between RS.MIA and MIA), 1 gene (*Ncmap, **fig. S7B***) showed exacerbated expression in response to the RS.MIA combination compared to control, and 1 gene (*Nes, **fig. S7B***) showed reduced expression (***Fig. 5D1***). 32 of the DEGs-RS.MIA_ipsi_ were also modified by RS alone (3.0% DEGs-RS) (***Fig. 5D2***). In contrast, on the contralateral side, only 28 of the 2094 annotated genes (1.3%) were also modulated by MIA in non-RS animals (***Fig. 5D3***); 26 were modulated to the same extent, while 2 genes (*Esrrg* and *Glmn, **fig. S7B***) showed exacerbated expression following RS exposure. 139 contralateral genes modulated by RS + MIA exposure were also affected by RS alone (6.7% DEGs-RS).

Importantly, pre-exposure to RS pre-empted MIA-induced changes for 119 DEGs-MIA, 75 on the ipsi side (***Fig. 5D1***, ***fig. S7C***, 75 of 693 = 11%) and 44 on the contralateral side (***Fig. 5D3***, ***fig. S7D***, 44 of 191= 23%), with 2 DEGs in common between both sides (*Srarp*, *Kcnh3***, *fig. S7C, D***). We focused on nociceptive signalling related genes and identified 14 DEGs on the ipsilateral side and 8 on the contralateral side. While RS exposure pre-empted MIA-induced anti-nociceptive transcriptional changes on the ipsilateral side (9 out of 14 reversed nociceptive DEGs, ***Fig. 5E***), it mainly mitigated the pro-nociceptive effects of MIA on the contralateral side (7 out of 8 DEGs, ***Fig. 5F***) suggesting different outcomes on nociceptive processing between the ipsilateral and contralateral side.

We also investigated DEGs only differentially regulated in the RS + MIA group, *i.e.* genes that were not regulated by RS or MIA exposure alone but by their combination only. 2,986 annotated genes were regulated by MIA only if animals had been exposed to RS prior to the MIA injection (920 on the ipsi side and 2066 on the contra side, ***Fig. 5D1****,**3***). 1,529 of those DEGs (856 ipsi and 673 contra) were no different between RS.MIA and MIA, suggesting no substantial impact of RS on the expression of these genes. However, 1457 (64 ipsi and 1393 contra, with 5 DEGs in common; ***fig. S7E***) of these were DEGs that were also differentially regulated between RS.MIA and MIA, suggesting a substantial impact of the RS.MIA combination on the expression these genes.

On the ipsilateral side (n=64), no remarkable pathways seemed enriched, with the exception of a couple of DEGs highlighted as being part the “Longevity regulating pathway”, (Enrichr, nominal *P*=0.044, ***Table S5***). However, on the contralateral side (n=1393), the prior exposure to RS led to major changes in a number of pathways (***Table S6***). Key regulators of longevity signalling were particularly affected (“Longevity regulating pathway”, nominal *P* = 0.000225; ***Fig. 6A***). Downregulation of *Foxo3*, *Stk11* and *Tsc2*, combined with upregulation of *Rheb*, *Rps6kb1* and *Eif4e*, suggested enhanced mTOR signalling, promoting anabolic processes while reducing stress resistance and autophagy^28,29^. Exposure to RS+MIA also led to changes in GABAergic signalling (“GABAergic synapse”, nominal *P* = 0.0205; ***Fig. 6B***) and glutamatergic signalling pathways (“Glutamatergic synapse”, nominal *P* = 0.054; ***Fig. 6C***). Overall, RS.MIA exposure appeared to enhance GABAergic inhibition through upregulation of GABA-A receptor subunits and Gng11, potentially strengthening inhibitory GPCR signalling. Glutamatergic signalling showed mixed effects: increased AMPA receptor expression suggested heightened excitatory transmission and plasticity, while reduced expression of NMDA/kainate receptor components, signalling enzymes, and scaffolding proteins indicated disrupted excitatory signalling and synaptic integrity.

**Figure 6.**
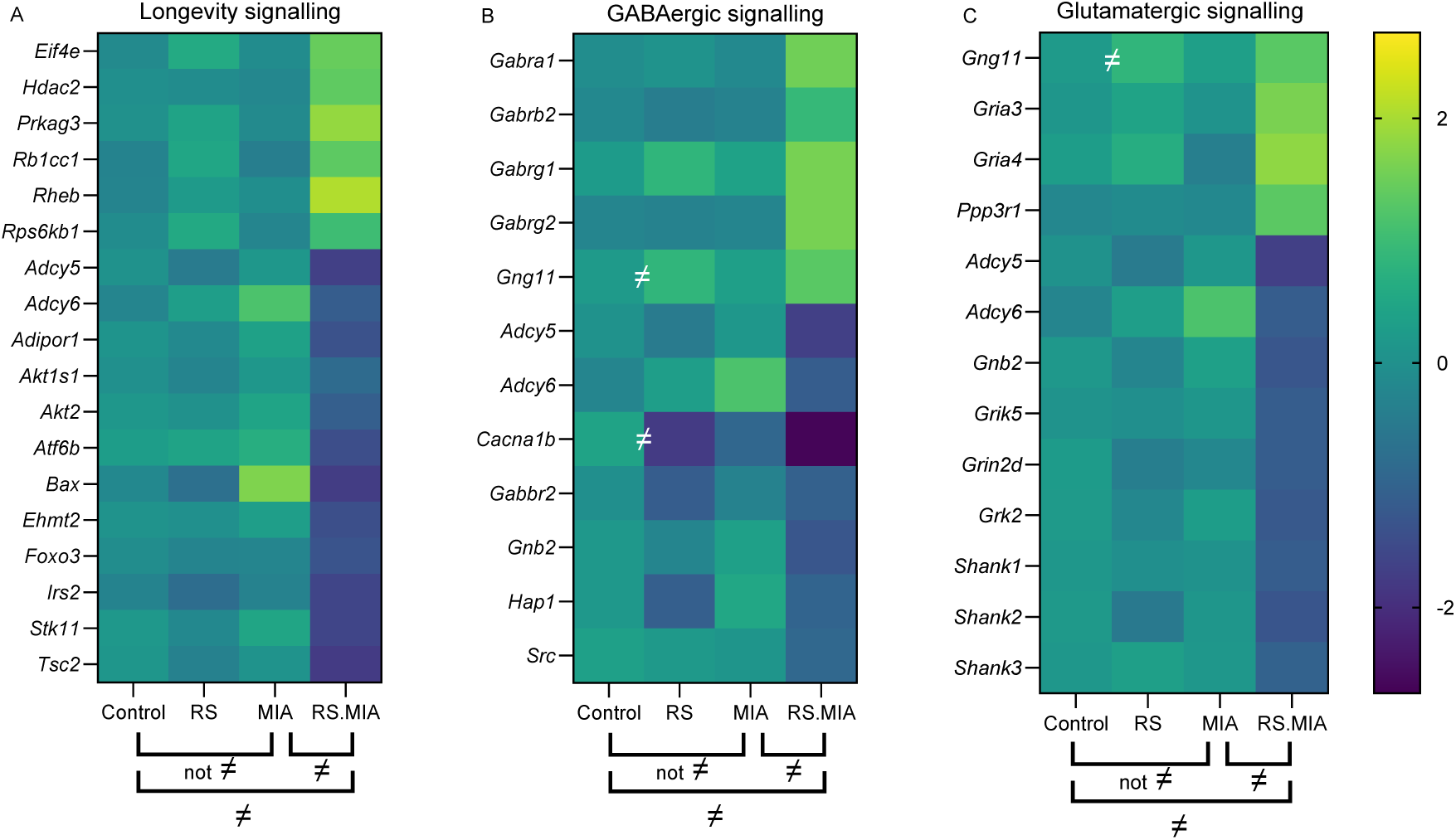
The combination of RS and MIA elicits unique signalling pathways absent in MIA-only animals. (**A–C**) Heatmaps showing differential expression (log₂ fold change) of genes involved in longevity signalling (**A**), GABAergic signalling (**B**), and glutamatergic signalling (**C**) across four experimental conditions: control, RS, MIA, and RS+MIA. Significant changes in gene expression between groups are indicated by “≠” brackets below the heatmaps. “≠” within heatmaps highlight genes with notable differential expression between Control and RS.

Overall, these results suggested that prior RS had 2 major outcomes after MIA: the reversal of some of the MIA-induced changes and more importantly the initiation of new gene programmes, with significant differences in modulation between the ipsilateral and contralateral side.

## 3. Discussion

How pre-existing individual characteristics, shaped by prior experience such as stress, influence the trajectory of pain throughout OA progression has received little attention. We demonstrate for the first time that exposure to stress can modulate subsequent pain outcomes in an established mouse model of OA in a sex-dependent manner. Importantly, our findings reveal that this evolutionarily conserved stress response can serve an adaptive role in the context of persistent pain. In male mice, RS attenuated MIA-induced pain- and anxiety-like behaviours and mitigated molecular changes in CNS regions implicated in stress and pain processing. In contrast, prior RS exposure exacerbated the pain experience in females, with no significant alterations observed at the molecular level. These results suggest that subchronic exposures to predictable stress in adulthood can influence the subsequent OA pain experience in a sex-dependent manner.

At the behavioural level, MIA injection led to the development of sensory and functional deficits as previously reported^30–33^. Both male and female mice displayed reduced mechanical withdrawal thresholds at the ipsilateral hindpaw, in agreement with previous reports ^32–35^. Interestingly, both sexes also developed mechanical hypersensitivity at the contralateral hindpaw, a phenomenon known as mirror-image pain (MIP), which resembles the symmetrical presentation commonly observed in human arthritic conditions^36–38^. MIP has been widely documented in various preclinical pain models^38–40^, but less commonly reported in models of osteoarthritis, probably reflecting historical emphasis on the ipsilateral side together with differences in experimental design including species and dosage of arthritis inducing agent^33,35,36,42–44^. The MIA injection also led to pronounced changes in weight bearing and dynamic gait behaviour, changes that were more prominent in the early phase of the joint disease across sexes. The assessment of pain and functional impairment performed with the CatWalk system further suggested a more prominent impact of knee injury in male mice than in females, as only MIA-injected male mice showed pronounced compensatory changes in gait. The sexually dimorphic effect of MIA may be further reflected in the weight of mice at the time of euthanasia as MIA attenuated body weight gain compared with controls in male mice only. The impact of MIA on emotional behaviour at one month was also more evident in males, with a significant decrease in the distance travelled and the probability of spending time in the centre of the open field arena, indicating an increase in anxiety-like behaviour. No parameters reached statistical significance in female mice, and this may be due to the early time point of assessment, as more robust displays of anxiety-like behaviour after MIA at later time points. Lastly, significant changes in adult hippocampal neurogenesis and GR expression within the PVN were observed in male, but not female, mice at one month after MIA injection. Together, these findings indicate that MIA alone had a more pronounced impact on both behavioural and molecular outcomes in male mice compared to females.

These sex differences were unexpected considering the clinical literature that often reports that women are more likely to develop chronic OA pain, experience greater pain for the same radiographic OA score, and present greater functional deficits than men^44,45^. However, our findings following the combination of subchronic restraint stress and MIA may come in agreement with the clinical observations, as females that had been exposed to stress showed greater hypersensitivity and anxiety-like behaviour. Specifically, RS exposure exacerbated the MIA-induced mechanical hypersensitivity in female mice while it attenuated this response in males. This bidirectional modulation of injury responses by stress was further reflected in some of the gait parameters at the time of peak functional deficit, as well as in anxiety-like behaviour assessed during a period marked by significant histopathological changes in knee joint architecture in this OA model^35,46^. Specifically, RS promoted the emergence of anxiety-like behaviours in females, but mitigated the MIA-induced anxiety-phenotype in males, suggesting that RS exposure predisposed females to a more vulnerable-like behavioural state in the context of OA-like joint pathology, while promoting a resilient-like phenotype in male mice. The significant correlation between mechanical sensitivity and anxiety-like behaviours further reinforced the dynamic interplay between sensory and affective components of OA pain and strengthen the clinical translatability of our findings, as pain severity has been shown to correlate with anxiety and depression symptoms in OA patients^15,47^. Of note, the correlation was stronger for measurements obtained from the contralateral hindpaw than from the ipsilateral hindpaw. Measurements at the ipsilateral hindpaw were taken closer to the site of injury and therefore likely to be confounded by primary hyperalgesia, whereas measurements at a distant site provided a more selective measure of centrally driven mechanisms. Our hypothesis that these mechanisms were likely to contribute to the resilient-like phenotype were also supported by the reduced second phase in the formalin test in stressed mice. In agreement with this, unilateral knee MIA injection was found to promote MIP development only in high-anxiety Wistar Kyoto rats, but not normo-anxiety Sprague Dawley male rats^32,48^. Our results highlight the modulatory role of prior stress in shaping sex-dependent susceptibility to MIA-induced outcomes, promoting more translationally relevant phenotypes in the context of OA pain and associated affective comorbidities.

Molecular analyses conducted one month after the MIA injection highlighted the mitigating impact of prior RS exposure on the behavioural response to MIA in male mice. Analysis of adult hippocampal neurogenesis revealed that the decrease in number of DCX positive neurons in male mice, as previously observed^30^, was prevented by prior RS exposure. This preservation of neurogenesis suggests that predictable, subchronic stress exposures may enhance hippocampal-dependent neurobehavioural flexibility, promoting adaptive responses to subsequent injury or environmental demands^49,50^. In parallel, RS prevented the MIA-induced increase in GR expression in the PVN of MIA-injected males, suggesting a preservation of neuroendocrine stress response regulation. These findings imply that in males, RS may mitigate the injury-induced behavioural impairments through establishment of an allostatic metastability at the molecular level in the CNS, supporting adaptive plasticity in response to sustained environmental challenges.

We next decided to continue our mechanistic analysis in male tissue only to focus on the novel, stress-associated, resilience-like phenotype observed in this sex. The results of an investigation of neuronal activity shortly after the MIA injection reinforced the notion of a stress-induced resilient-like phenotype in males. Prior exposure to subchronic stress normalised the injury-induced upregulation of c-Fos expression in the deeper layers of the spinal dorsal horn, as well as in key supraspinal regions involved in pain modulation and stress integration such as the PAG and PVN. Interestingly, we found no differences in c-Fos expression in the superficial laminae of the dorsal horn or in the PBN, first relay distributing spinal nociceptive signals to forebrain areas. These observations suggested that prior stress exposure did not alter primary nociceptive input, but rather influenced higher-order supraspinal contributions to pain processing, such as descending modulatory pathways, which are known to shape contralateral pain^51–56^. This may explain why the impact of RS on mechanical threshold were more pronounced on the contralateral than on the ipsilateral side. RNAseq analysis conducted at one month after the MIA injection provided further insight into the molecular changes supporting the resilient-like phenotype in males. Prior restraint stress reprogrammed MIA-induced transcriptional changes at spinal cord level. Notably, RS pre-empted a subset of these changes, altering nociceptive gene expression in a side-specific manner. Crucially, RS reversed more pro-nociceptive changes on the contralateral side than on the ipsilateral side, which is again consistent with the more pronounced behavioural effects of RS observed contralaterally. Prior RS also triggered novel transcriptional programs upon MIA injection. RS followed by injury promoted a strong compensatory upregulation of GABA-related genes – suggesting a homeostatic rebound in inhibitory tone – which was more apparent on the contralateral side. Upregulation of *Gabra1*, *Gabrb2*, *Gabrg1*, and *Gabrg2* suggested enhanced fast synaptic inhibition via GABA-A receptors, likely leading to reduced neuronal excitability. Although the transcriptomic profile revealed both enhanced excitatory and inhibitory signalling components, the net outcome of reduced hypersensitivity suggests that the strengthening of GABAergic inhibition dominates the functional landscape. This interpretation is supported by the significant downregulation in RS+MIA mice of *Src,* known to have a major role in enhancing excitatory transmission through NMDA receptor phosphorylation in the context of persistent pain^57^. Overall, stress preconditioning appears to have reprogrammed the injury response, priming neural circuits to mount an exaggerated inhibitory (GABAergic) adaptation after injury, which may reflect a form of homeostatic plasticity. This pattern supports a model of stress-primed neuroplasticity, where prior adversity can trigger adaptive responses to subsequent challenges, rather than purely maladaptive ones.

However, the combination of RS and MIA also disrupted transcriptional programs linked to cellular longevity suggesting long-term costs of the double exposure to trauma. The gene expression profile reflected a shift away from protective, longevity-associated programs toward a metabolically active, stress-sensitive state. In post-mitotic cells such as neurons, this trade-off may heighten vulnerability to aging and degeneration by increasing demand while weakening long-term cellular resilience^58,59^.

In summary, we demonstrate that prior stress exposure can influence the pain experience in a mouse model of OA in a sex dependent manner. While subchronic restraint stress exacerbated pain- and anxiety-like behaviours in female mice following joint injury, it attenuated these outcomes in males, promoting a resilient phenotype. Whether the differential impact of RS on males and females is linked to sex-specific responses to MIA has not been explored in this study but remains a possibility that should be investigated. The resilience in males was accompanied by the prevention of several MIA-induced changes in molecular markers within limbic brain regions and at spinal level. However, it also coincided with transcriptional alterations in the spinal cord suggestive of shifts in cellular function and stress resistance, raising the possibility that such resilience may be accompanied by downstream costs following injury. Of note, although the present study identifies stress-sensitive neural and molecular signatures associated with a resilient-like phenotype, causal testing of these pathways will require targeted circuit- and molecular-level manipulations in future work. Importantly, resilience should not be viewed as a uniformly beneficial molecular or behavioural state but instead as an emergent outcome of complex interactions with the environment, whereby adaptations in pain salience, cognitive–evaluative processing and motivational drive shape pain expression. In this context, our findings advance understanding of the biological heterogeneity underlying chronic pain and highlight the modulatory role of the environment in conferring vulnerability or resilience to persistent pain states.

## 4. Methods

### 4.1 Animals and housing conditions

Male and female C57Bl/6J mice were supplied by Charles River UK at 8 weeks of age. Female mice were freely cycling and randomly assigned to groups; under this design, any effects of oestrous cycle stage are expected to average out and not systematically confound group comparisons^60–63^. Upon arrival at the Central BSU facility, mice were randomly housed in groups of 3-5 in a temperature (22±2°C) and relative humidity (50±10%) controlled environment, under a 12h light/dark cycle, with gradual lights on/off at 7am/7pm, respectively, and provided with food (Teklad Global 2018 diet) and water *ad libitum*. Cages were fitted with sawdust bedding and comfort fresh nesting materials (Datesand Cocoons), a cardboard tunnel, and several chewstick blocks. Animals were allowed a minimum of one week acclimatization before the start of experiments and were handled via tunnel when possible. All procedures were carried out under the Home Office License P8F6ECC28 in accordance with the guidelines of the UK Animals (Scientific Procedures) Act 1989 and subsequent amendments.

### 4.2 Study design

Animal numbers were determined using power calculations based on our previous studies^41,64^. A total of 102 mice were allocated to 5 studies as follows: a| behavioural and molecular characterization of the effect of RS on the MIA-induced pain experience included two separately ran cohorts of animals consisting of n = 22 male and n = 24 female mice (experiment started with a larger female cohort to account for the expected higher mortality rate occurring in this sex after the MIA injection); b| n = 12 male mice were used to investigate the impact of RS on hippocampal neurogenesis; c| n = 20 male mice were used to characterise the impact of RS on spontaneous nocifensive behaviours after formalin; and d| fresh spinal cord tissue of n = 24 male mice was used for the RNA sequencing study. For all studies, the intervention (stress/injury) was performed per cage to prevent possible changes in the sensory and/or affective phenotypes of non-exposed animals.

All behavioural testing was conducted during the animal’s inactive phase, with all mice tested at the same time of the day. Throughout the experiments, we aimed to reduce the impact of stress that could result from multiple testing sessions and to increase the habituation of mice to the experimental routine whilst also limiting prolong disruptions to their sleep. All behavioural experiments were carried out between 7:30am to 12pm. The only exception to this schedule was the novel object recognition test, due to its habituation and testing timeline.

Because the MIA model induces visible gait abnormalities and guarding behaviour especially in the first 1-2 weeks after the i.a. injection, full experimental blinding for injury status could not be maintained during testing. However, all behavioural and molecular analyses for the stress condition were performed with the investigator blinded to treatment group to ensure unbiased data collection and analysis. Unblinding occurred after statistical analysis.

### 4.3 Subchronic restraint stress

To induce stress, mice were restraint for 1h by being placed in 50ml falcon tubes fitted with air holes and a stopper to allow them to breathe, but prevent their escape, as previously reported^65,66^. This procedure was carried out over three consecutive days, starting each time at 8:30am, with mice being given no time to habituate to the room. Non-stressed mice were left undisturbed in their home cages throughout this time.

### 4.4 MIA injection

To induce knee osteoarthritis, monoiodoacetate (MIA) was intra-articularly injected as previously reported^30,67^. Specifically, mice were placed under general anaesthesia using 2% isoflurane in O_2_ (flow rate 1.5 L/min). After cessation of reflexes, the mouse was placed on its back and the area surrounding the left knee was trimmed and wiped with alcohol. The knee was stabilised in a 90° bent position, and the white patellar tendon region just beneath the patella was lightly marked by applying gentle pressure on the skin. Freshly prepared monoiodoacetate (MIA; 10μl of 10% w/v MIA, Sigma-Aldrich, I2512, dissolved in sterile 0.9% NaCl; as widely used in pain research labs^41,67^) solution was injected, perpendicular to the tibia, through the marked knee joint area. After needle retraction, the hindleg was gently moved back and forth to ensure even distribution of the solution. Due to the high toxicity of the compound, the health status of mice was closely monitored in the next 72h.

Control mice did not receive an i.a. injection of saline to prevent any potential damage to the knee joint, including the patellar and cruciate ligaments, menisci, and possible inflammation of the synovial membrane, that could result from this procedure and would confound the results.

### 4.5 Formalin injection and nocifensive behaviour

The impact of RS on the pain experience was also assessed using the subcutaneous model of inflammatory/chemical pain induced by formalin (methods adapted from ^68^). To limit potential stress-related confounds that could influence the nocifensive behaviour, mice (*N* = 20) underwent a series of habituation sessions to the behavioural room, experimental setup, and experimenter, both before and after exposure to the RS. Two habituation sessions lasting 1h each were conducted one day apart before the RS exposure and they consisted of placing the mice in opaque Plexiglass chambers (9.5cm length, 14cm height) positioned on an elevated wire grid (Ugo Basile, Italy). Three days later, *N* = 10 mice were exposed to the restraint stress protocol. Following the final RS session, two additional habituation sessions were carried out at 2-4-day intervals. Two weeks after the start of restraint stress, mice were returned to the behavioural room for assessment of nocifensive behaviours.

On the test day, mice were acclimated to the experimental room for 1h before being placed in a restraining tube and administered a subcutaneous (intraplantar) injection of 10μl of 5% Formalin (Sigma-Aldrich) diluted in 0.9% NaCl (B Braun) into the left hindpaw. The injection site was wiped dry when needed, and mice were placed in the plexiglass chambers, with their behaviour recorded for 60min using a camera placed underneath the setup. If mice fell asleep at any time during the recording phase, they were left undisturbed. At the end of the 60min time period, mice were returned to a new cage to prevent stressing their uninjured cage-mates still present the original transport cage.

At 2h after formalin injection, mice were placed under terminal anaesthesia and perfused transcardially for tissue collection. Videos acquired using the EthoVision XT14 (Noldus Information Technology) software were analysed blindly using the software’s manual tracking function. Licking and flinching behaviours of the injected paw were analysed in 5min bins. The first phase of the nocifensive behaviour was defined as the time from 0 - 10min post injection, while the second phase as 10.01 - 60min.

### 4.6 Mechanical sensitivity

The static mechanical withdrawal threshold was assessed using the up-down method as previously described by Chaplan et al.^69^ and previously applied in our lab^30^. Mice were individually placed in opaque Plexiglass cubicles (9.5 x 9.5cm, 14cm h) situated on an elevated metal mesh stand (Ugo Basile) and allowed to habituate the enclosure and experimenter’s presence for as long as necessary, but no longer than 3h. A series of calibrated von Frey monofilaments (Ugo Basile) were used to determine the withdrawal threshold at the plantar surface of the hindleg. Testing began with filament 3.61 (weight 0.4g). A positive response – defined as a brisk paw withdrawal, splaying of the digits, or otherwise the mouse directing its attention to the just stimulated hindpaw - led to the application of the next lower force filament in the series (3.22, weight 0.16g) after a minimum 3min interval. In the absence of a response, the next higher force filament was applied. After the first change in response pattern, four additional measurements were taken and the paw withdrawal threshold (PWT) was calculated using the formula: 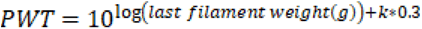, where *k* is a constant value corresponding to each individual pattern obtained^70^.

### 4.7 Static weight bearing

Static weight distribution between the hindlegs was assessed using the Bioseb Incapacitance Test (Bio Swb-Touch-M), as previously reported^30^. To ensure adequate compliance with the task, mice underwent approx. six sessions of habituation and training to familiarise them with the experimenter, apparatus, and the task. Before testing, mice were allowed to acclimate to the testing room for at least 30min. Mice were then placed inside the apparatus where they were required to maintain a stable, upright posture for at least 7sec. Up to 5 measurements were collected per session when mice complied with the testing. The values obtained were then averaged and the weight distribution was calculated using the formula: 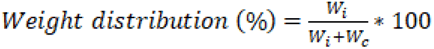, where W_i_ and W_c_ are the weights in grams borne on the ipsi-and contralateral hindlegs, respectively.

### 2.7 Dynamic gait analysis

Objective quantification of unidirectional locomotion was performed using the CatWalk^®^ eXtended Technology system (Noldus Information Technology). The apparatus consists of a 1.3m-long corridor with an internally LED illuminated glass walkway. As the mouse spontaneously transverses the corridor to reach a safe space located at the opposite end of the walkway, contact between the paws and the glass surface scatters the green LED light within the walkway. This scattering is captured by a high-speed camera placed beneath the walkway. A red LED light illuminating the corridor form the ceiling creates an optimal contrast to capture the animal’s footprints. Given that the testing occurred during mice’s inactive phase, animals were not given a time to habituate to the behavioural room, to prevent their unwillingness to run during testing. A run was considered compliant when the speed of the mouse was between 17 – 30 cm/s, with a maximum variation below 60%. The green intensity threshold (0.10) and camera gain (19.58) were constant throughout the experiments for inter-animal paw comparison. A minimum of two compliant runs were included in the analysis, and the parameters assessed were based on previous literature on arthritis models^31^.

### 2.8 Open field test

Anxiety-like behaviour was assessed using the open field test. The apparatus consisted of a square arena measuring 90 x 90cm, with 40cm high walls, uniformly illuminated from above it. The illumination in the centre of the Open Field was 500 lux. Mice were allowed to acclimatise to the room for 40min before being placed, individually, in the center of the arena. Their movements were tracked using the EthoVision XT14 software for 5min using a camera positioned directly above the apparatus. After each trial, the arena was cleaned with 70% ethanol to eliminate olfactory cues.

Raw movement data (x-y coordinates and movement velocity) were exported and analysed using a Python script. The geometric center of the arena was determined as the midpoint between the minimum and maximum x and y coordinates recorded during the trial. Using this midpoint, three concentric circular zones were defined: ‘Near Center’ zone (within a 20 cm radius, R ≤ 20 cm), ‘Away from Center’ zone (20 cm < R ≤ 40 cm), and ‘Far Away from Center’ zone (40 cm < R ≤ 60 cm). For each frame, the Euclidean distance between the animal’s position and the arena midpoint was calculated to classify the position into one of these zones. The proportion of frames in each zone was computed to quantify the probability of the animal occupying the three zones of the open field arena during the trial.

Traveling velocity values, exported from EthoVision, were averaged for each defined zone to indicate the animal’s movement pattern. Total distance moved in each zone was calculated by summing the distance moved every 0.2sec within the three zones. To determine the probability of the mouse being located in a specific zone, a Boolean array mask was created where each element was either True (1) if the mouse center point fell within the respective zone, or False (0) otherwise. The mean value of each mask array represented the probability for each respective zone. Representative images were created using the scatter function from Matplotlib to plot mouse center points that fall within each mask, using different colours for each zone.

### 4.8 Novel object recognition

To quantify novel preference, the novel object recognition test was employed as previously described^30^. After a 30min habituation period in the experimental room, each mouse was placed, wall-facing at the far-end of a circular arena (diameter 35cm, height 30cm), opposite to the location of two identical objects (circular glass bottles: diameter 6.5cm, height 17cm, brown in colour; denoted as ‘familiar’ objects). Mice were allowed to freely explore the enclosure for 10min, and become familiar with both object and the arena, with their movements being continuously tracked using a camera connected to EthoVision XT14 software. After this ‘habituation/familiarisation’ phase, establishing familiarity with the two identical objects, mice were returned to their cages while the arena and the objects were thoroughly wiped with 70% ethanol. Four hours later, animals were placed back inside the arena, now containing one of the original (now ‘familiar’) object, and one novel object (elliptical glass bottle: length 8.5cm, width 4.5, height 15cm, white in colour). During this ‘test’ phase, the mouse location was recorded for an additional 5min. The position of the novel object was alternated relative to the familiar object between trials to account for possible side preferences.

Object exploration was considered when the mouse’s nose-point was within a 2cm distance from either object. The amount of time spent exploring the objects was calculated using the formula: 𝐸𝑥𝑝𝑙𝑜𝑟𝑎𝑡𝑖𝑜𝑛 of novel (%) = 𝑡_𝑁_/(𝑡_𝑁_+𝑡) ∗ 100, where t_N_ and t_F_ is the time spent exploring the novel and familiar object, respectively. For the habituation/familiarisation phase, the ‘novel’ object was treated as the familiar one that would later be replaced with the novel object during the testing phase.

### 4.9 Tissue fixation and immunofluorescence

At one month (30 days) after the i.a. injection of MIA or day 12 for RS only and Formalin group, mice were placed under terminal anaesthesia using a gas mixture of isoflurane (2% mixed with O_2_, flow rate 1.5L/min) for induction, followed by an intraperitoneal administration of pentobarbital (Dolethal) overdose. After cessation of reflexes, mice were transcardially perfused with 20ml NaCl solution containing Heparin (5000IU/ml), followed by 20ml of ice-cold 10% formalin solution (Sigma-Aldrich, HT-501640-19L). Brain and spinal cord tissue was dissected and postfixed in 10% formalin solution for an additional 2h at 4°C, then placed in 30% sucrose solution containing 0.5mg/mL NaN_3_ for a minimum of 3 days, or until they sank. Tissue was then coronally sectioned on a freezing microtome (Leica) at 40µm thickness and free-floating sections were placed in 5% sucrose containing 0.5mg/mL NaN_3_ and stored at 4°C until further processing.

For c-Fos and GR staining, free-floating sections were first rinsed with 0.1M PB for 10min then incubated with 30% normal donkey serum (Biosynth, 88R-D001) in 0.1M PB containing 0.3% Triton X-100 for 1h to prevent nonspecific binding. Tissue was then incubated overnight at 18°C with either guinea pig anti-c-Fos (1:2000, Synaptic Systems, 226308) or mouse anti-GR (1:500, Santa Cruz, sc-393232). After 3 x 10min washes in 0.1M PB, sections were incubated with respective secondary antibodies diluted in TTBS (anti-guinea pig Alexa 594, Jackson ImmunoResearch, 706-585-148; anti-mouse Cy3, Jackson ImmunoResearch, 715-165-150) for 2h in the dark. Sections were washed one last time for a total of 30min then mounted on gelatinised slides and coverslipped with Fluoromount Aqueous Mounting Medium (Sigma-Aldrich, F4680), and stored at 4°C.

Adult hippocampal neurogenesis was detected using a primary antibody against doublecortin (rabbit anti-DCX, 1:5000, Abcam, ab18723). After overnight incubation at 18°C, sections were washed 3 x 10min in 0.1M PB, then incubated for 2h with anti-rabbit biotinylated secondary antibody (1:400, Bethyl Labs, A120-108B). After another round of washes, tissue was incubated with preformed avidin-biotin-HRP complexes (1:200; Vectastain ABCKit Peroxidase, PK-6100) for 30min. Finally, tissue was incubated for 5min with tyramide solution (Ape Bio, A8011-APE), washed 2 x 15min with 0.1M PB, then incubated for an additional 2h with Avidin D, Fluorescein labelled (Vector Labs, A-2001-5). Sections were washed, mounted on gelatinised slides and lastly coverslipped as described above.

### 4.10 Image acquisition and analysis

DCX immunopositive cells were manually counted in the dentate gyrus of the dorsal hippocampus (bregma –1.22mm to –2.3mm) using a Leica (Nussloch, Germany) DMR microscope. The presence of a DCX^+^ cell body, with or without an axonal projection through the granule cell layer of the dentate gyrus, was considered a positive cell. For c-Fos staining, images of lumbar spinal cord were taken using the Leica microscope connected to a Hamamatsu (C4742-95, Shizuoka, Japan) digital CCD camera and Velocity 6.3 software. Positive cells were manually counted in the dorsal horn of the spinal cord using the cell counter plugin in Fiji software.

For supraspinal immunofluorescence analysis of c-Fos and GR, images were acquired using a Zeiss AxioScan Z1 slide scanner coupled to Zen Microscopy software. Positive cells stained for c-Fos were automatically quantified using custom Fiji (ImageJ) macros. In the PAG (bregma –3.8 to –4.96mm), three background intensity measurements were taken in cell-free areas lacking visible staining, within the same subregion, across the *dm*, *l*, and *vl*. These values were averaged and multiplied by two to define the lower threshold for signal detection. Particle analysis was then performed (pixel^2^ = 10 – Infinity, circularity = 0.3 – 1.00) to quantify positive cells. For the PBn (bregma –5.02mm to –5.34mm), LC (bregma –5.34mm to –5.68mm), and PVN (bregma –0.7mm to –0.94mm) a single background intensity measurement was made in a representative cell-free area within the same region, and the same particle analysis parameters were applied. For GR quantification in the dorsal hippocampus, circular ROIs were used in the CA1-3 and DG regions to measure mean fluorescence intensity within the cellular layers. Background intensity was measured within the corresponding axonal/fiber zone adjacent to each cellular layer ROI and this value was subtracted from the respective ROI measurement prior to analysis. In the PVN and in the CeA and BLA amygdalar nuclei, the regions of interest were manually delineated for each section, and GR positive cells were then quantified using the analyse particles plugin (pixel^2^ = 7.5 – Infinity, circularity = 0.3 – 1.00), applying a threshold set at twice the measured background intensity. For cFos stain in the RVM, positive cells were counted directly under the microscope. For each stain, the total number of cells per region was computed, with minimum of 3 sections averaged per animal.

### 4.11 RNA sequencing

RNA was extracted from lumbar spinal cord dorsal horn quadrants and prepared as previously described^41^. mRNA libraries were prepared from total RNA using NEBNext Ultra II Directional Library Preparation Kit at the Genome Center. Queen Mary University, RNA sequencing data were analysed in R Studio version 4.2.2 or later using established pipelines. In brief, paired-end next generation sequencing reads (75 bp) were trimmed of low quality (Q > 20) bases and adapter sequences then aligned to the mm10 reference genome. Gene counts were quantified and filtered using a minimum count threshold of 10, then libraries were normalised by the TMM method. Gene expression changes were identified using linear models fitted to log-transformed counts per million (log-CPM) values with moderated variance estimates. Genes exceeding a 1.2-fold change in expression and nominal significance less than 0.05 criteria were considered differentially expressed and advanced for further downstream analysis, using both Ingenuity Pathway Analysis (IPA) software (Qiagen, Redwood City, CA, USA) and Enrichr (https://maayanlab.cloud/Enrichr/) with a focus on KEGG pathways, based on Fisher’s exact test and ranked by combined score pathway analysis. Cell-type proportions were estimated from bulk RNA sequencing using reference-based deconvolution against the mouse single-cell atlas^71^.

### 4.12 Statistical analysis

Data were analysed in GraphPad Prism (version 10.5.0 (774)) and IBM SPSS Statistics (version 331.0.0.0 (171)). Individual animals were considered the experimental unit. Group sizes were determined based on power calculations conducted for similar studies^30,41,64^. Statistical analysis was mostly performed using Student’s *t* test, 1-way or 2-way repeated measures (RM) ANOVA, as well as simple linear regression. Multiple comparisons were performed using Šídák’s method. Mixed effects analysis was used for data-sets with missing data-points. Normality was assessed using the Shapiro–Wilk test and inspection of Q–Q plots, which provide a more robust assessment when formal tests are overly sensitive. For von Frey thresholds, raw data were log-transformed to normalize distributions and stabilize variance. Residuals rarely deviated from normality and Q–Q plots showed no severe departures, supporting the validity of parametric analyses. Where significant deviations from normality occurred, corresponding non-parametric tests were also used. Significance was set at *P* < 0.05. When the assumption of sphericity was violated in RM ANOVA, the Greenhouse Geisser correction was applied. Data are presented as group means ± standard error of the mean (SEM), with the exception of box-and-whiskers plots, which display the IQR. Outliers were identified objectively using Tukey’s method, based on IQR, as indicated in ***Table S1,*** but data points removal was not automatic. For behavioural data, data points were excluded only when the same animal was an outlier in at least two outcome measures from a specific behavioural category (e.g. catwalk, open field parameters), in which case the animal was removed from all parameters related to this behavioural category. For molecular observations, data points were only excluded when blind and independent observations confirmed technical issue with the stain (*e.g.* poor perfusion, insufficient number of sections containing the area of interest).

As simultaneous testing male and female mice can introduce confounding variables (e.g., olfactory, auditory) that may influence behavioural outcomes, male and female cohorts were run independently and analysed separately. The statistical test used for each comparison is reported in the supplementary ***Table S1***; F and *P* values are reported in the main text, while the *N* values (sample sizes) are provided in the figure legends, with detail breakdowns in ***Table S1***. For supplementary figures, the statistical test used, as well as the F, *P*, and *N* values are indicated in the figure legends.

## Acknowledgements

This project was funded by a Versus Arthritis grant to SG, Research Award 21972. SS was funded by the Advanced Pain Discovery Platform (MR/W002566/1). LA was funded by a grant from the Medical Research foundation MRF-087-0001-F-KOCH-C0917 and CJB was funded by a Brain Research UK fellowship.

## Author contributions

SG and RF conceived the study and designed all experiments. RF conducted all behavioural and molecular experiments. SS ran all sequencing analysis. LA and CJB contributed to data analysis. SG and RF jointly analysed and interpreted the data and wrote the manuscript first draft. SH contributed to the establishment of the MIA model and behavioural assessment paradigms, and provided critical insights throughout the study. All authors provided critical comments and revisions on the draft and all authors read and approved the final manuscript.

---

**Figure S1:**
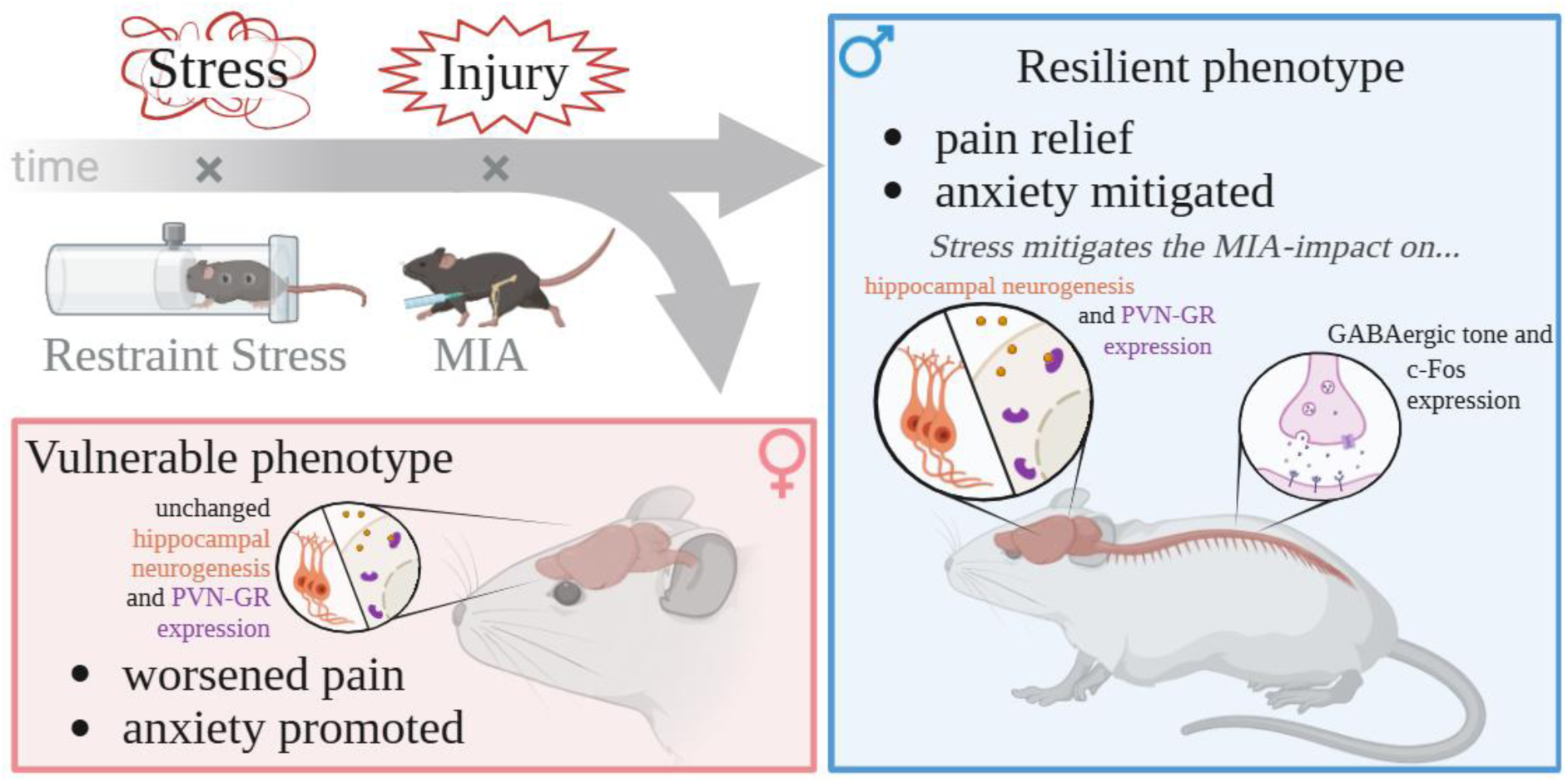
Graphical abstract: prior sub-chronic restraint stress modulates the long-term pain experience in osteoarthritis in a sex-specific manner. In male mice, stress promoted resilience, reducing MIA-induced mechanical hypersensitivity, gait deficits, and anxiety-like behaviour, while females showed mild symptom exacerbation. Stress reprogrammed spinal and brain injury responses in male mice, priming GABAergic compensation and dampening pain-related transcriptional changes.

**Figure S2.**
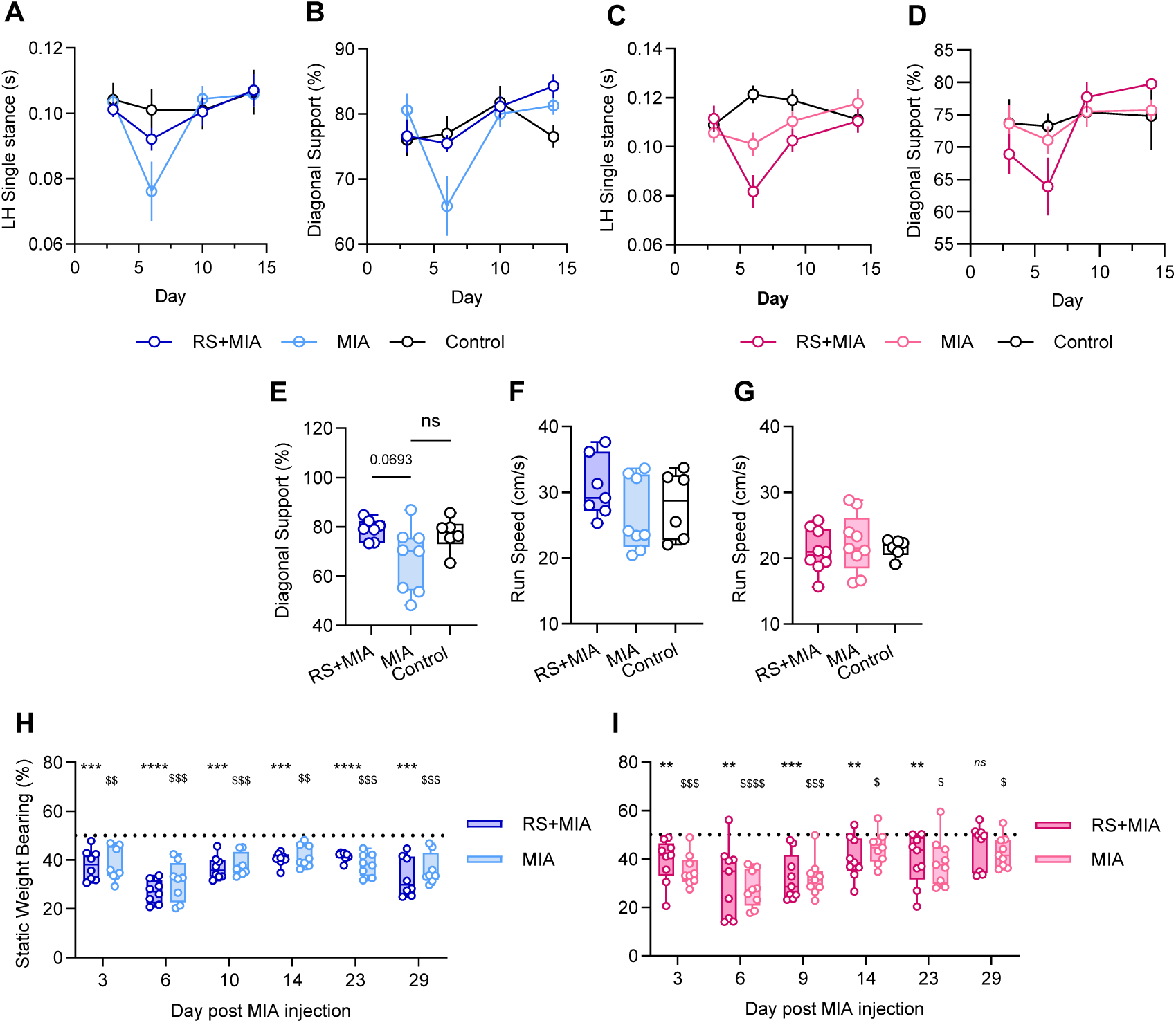
Time course of dynamic gait behaviours and static weight bearing after MIA injection. (**A**-**D**) Time course of selective gait parameters in male (**A**: mixed-effects ANOVA: ‘Time’ F_3,54_=9.15, *P* < 0.0001; **B**: mixed-effects ANOVA: ‘Time’ F_3,54_=8.815, *P* < 0.0001; *N* = 7/8/6) and female mice (**C**: mixed-effects ANOVA: ‘Time’ F_3,58_=4.74, *P* = 0.005; ‘Treatment’ F_2,21_=3.53, *P* = 0.0476; **D**: mixed-effects ANOVA ‘Time’ F_3,58_=6.66, *P* = 0.0006; *N* = 9/9/6). (**E**) Bar plot illustrating diagonal support during gait in male mice at day six after MIA injection (1-way ANOVA, F_2,18_=3.542, *P* = 0.05, Šídák’s post-hoc analysis). Bar plots illustrating the travelling speed of male (**F,** N=7/8/6) and female (**G,** N=9/9/6) mice at day six after the MIA injection. *N* = 9/9/6. (**H**, **I**) Static weight bearing distribution in male (**H**, *N* =8/8, \*\*\**P* < 0.001, \*\*\**P* < 0.0001, RS+MIA *vs* 50%; ^$$^*P* < 0.01, ^$$$^*P* < 0.001, MIA vs 50%) and female (**I**, *N* = 9/9, \*\**P* < 0.01, \*\*\**P* < 0.001 RS+MIA *vs* 50%. ^$^*P* < 0.05, ^$$$^*P* < 0.001, ^$$$$^*P* < 0.0001 MIA *vs* 50%.) mice. Nominal significance was computed by conducting separate one-sample t-tests for each group across days.

**Figure S3.**
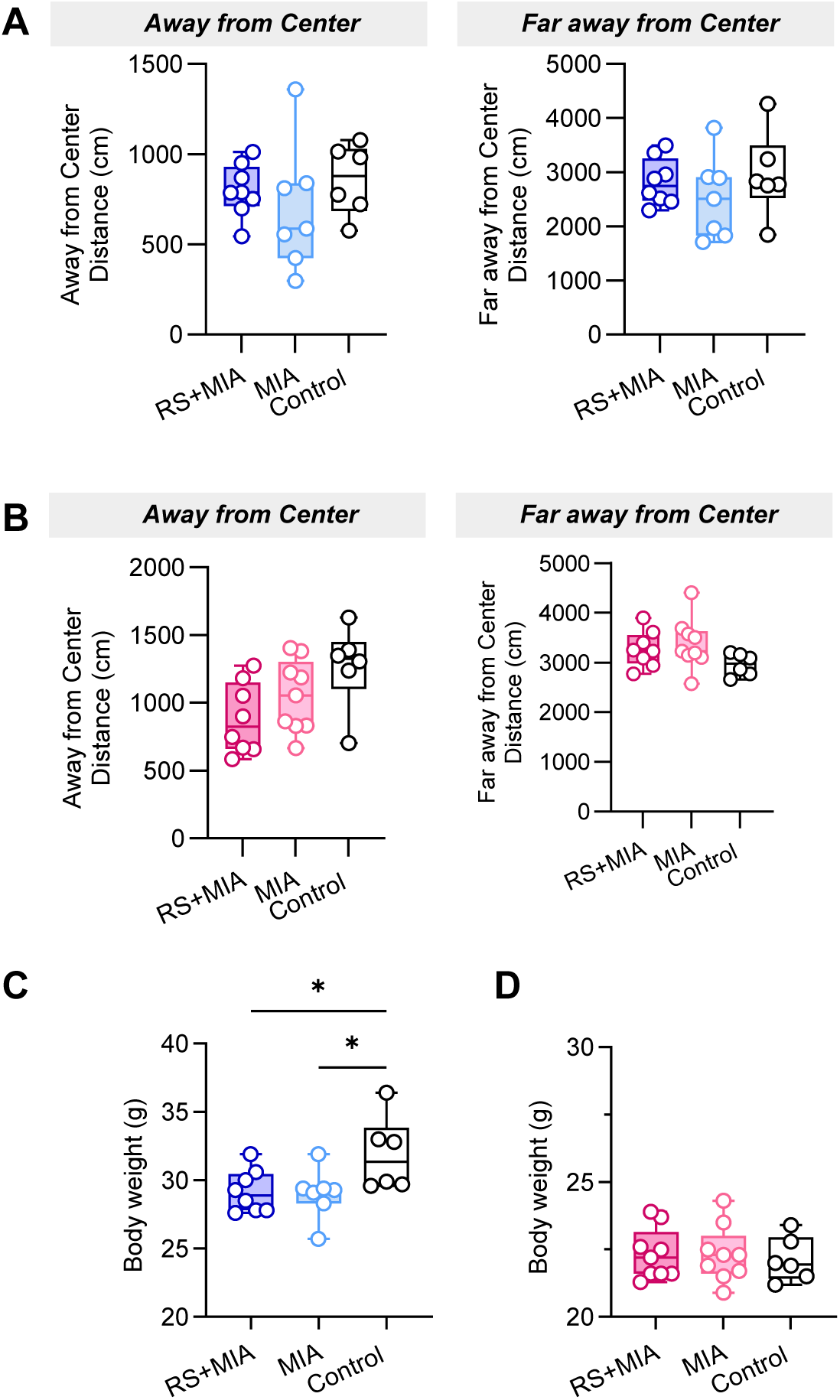
Extended characterization of anxiety-like behaviour, correlations with pain-like responses, and general health parameters. (**A**, **B**) Total distance travelled by male (**A,** N=8/7/6) and female (**B,** N=8/9/6) mice in the respective zones of the open field arena at day 29 after the MIA injection. (**C**, **D**) Body weight of male (**C**, 1-way ANOVA, F_2,18_=4.11, *P* = 0.0339; *N* = 8/8/6) and female (**D**, *N* = 9/9/6) mice at the time of euthanasia (day 30 after MIA injection).

**Figure S4.**
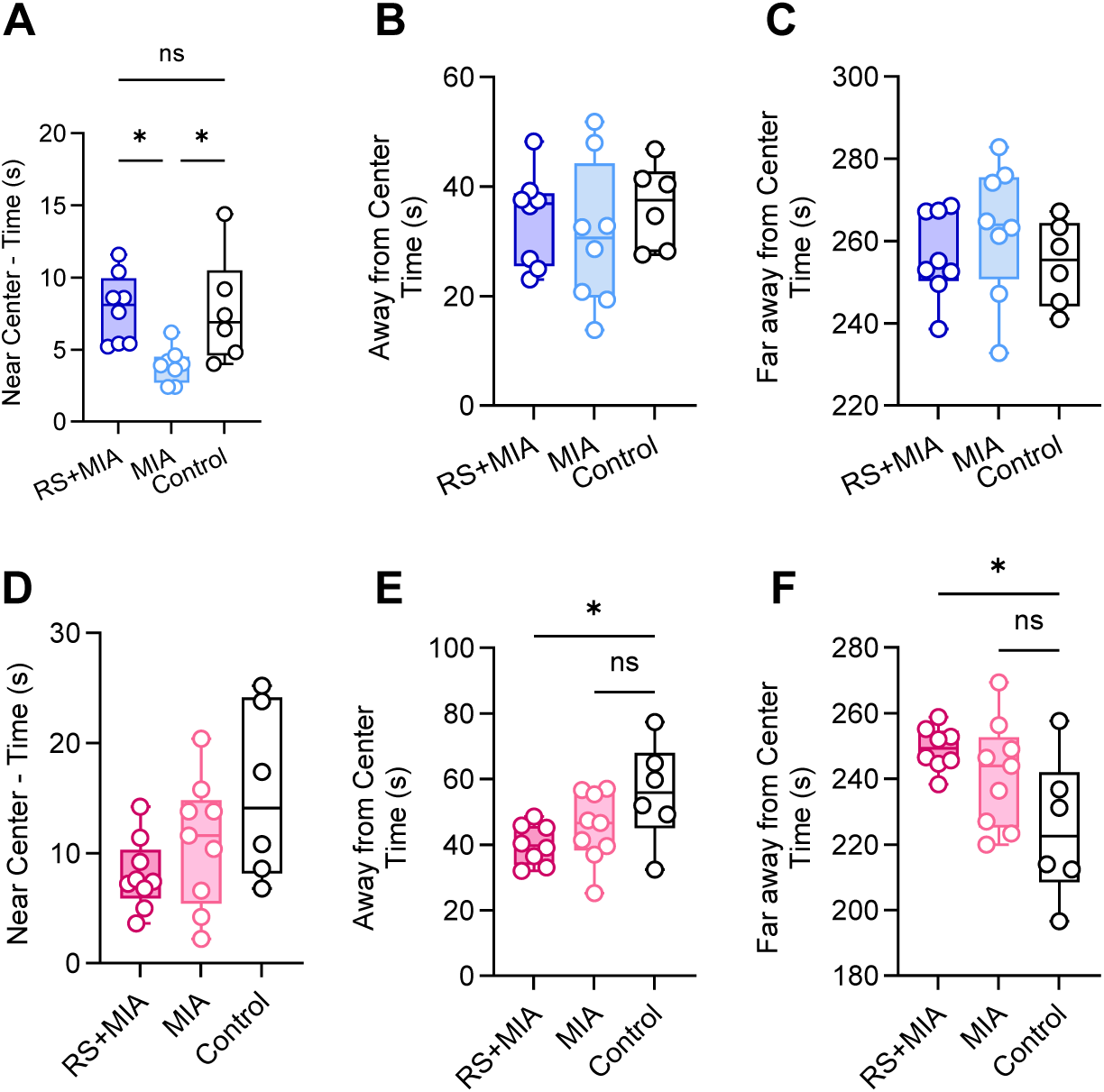
Characterization of anxiety-like behaviour using time spent in different zones of the arena. **(A)** F_2,19_=5.94, *P* = 0.0099. **(D)** F_2,21_=3.103, *P* = 0.066. (**E**) F_2,20_=3.792, *P* = 0.0401. (**F**) F_2,20_=4.407, *P* = 0.026. Šídák’s post-hoc. \**P* < 0.05. A-C: *N* = 8/8/6, D-F: *N* = 9/9/6.

**Figure S5.**
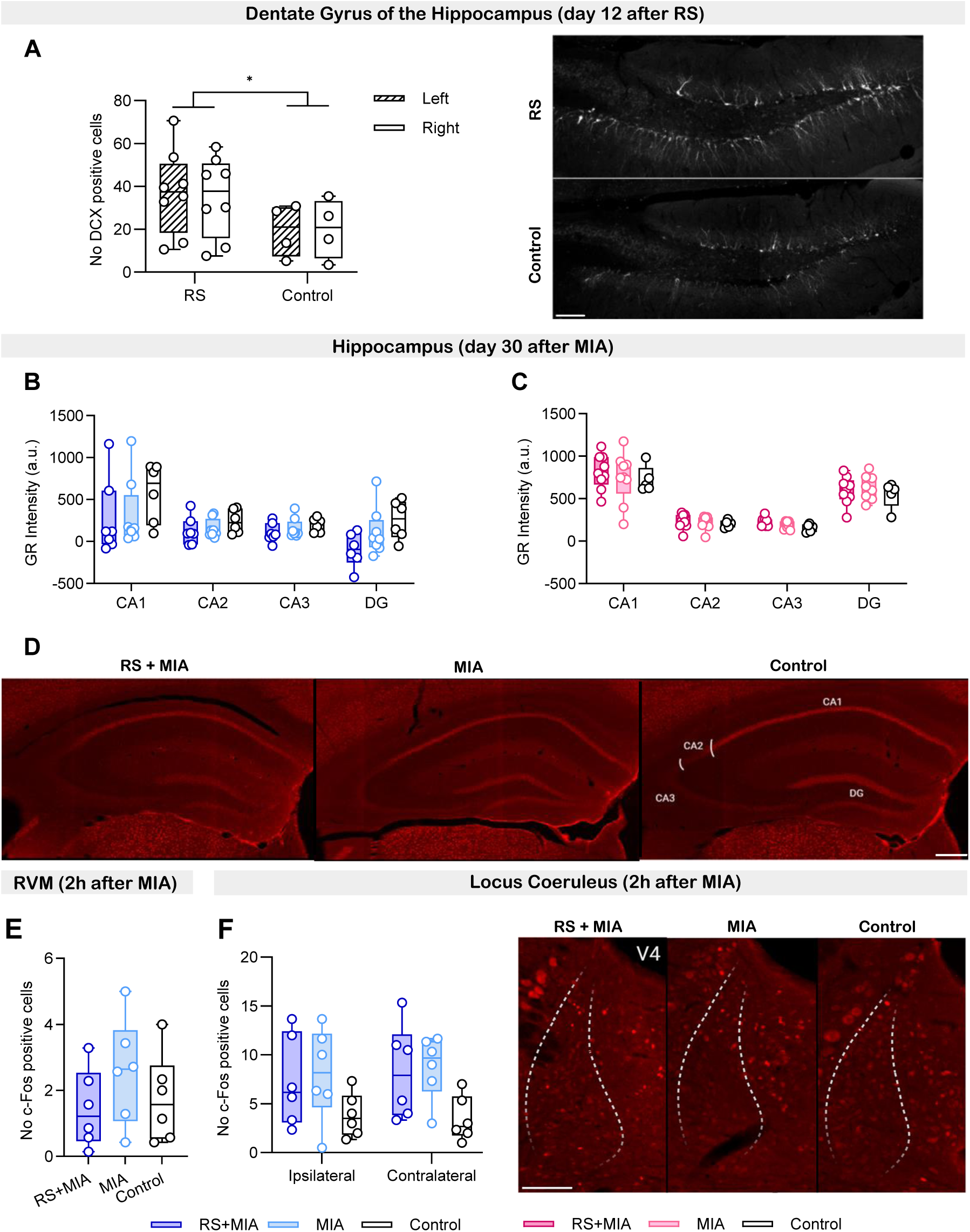
Molecular observations following exposure to RS, MIA or RS+MIA. **(A**) Quantification (left) and representative images (right) of DCX positive cells in the dentate gyrus of the hippocampus at two weeks after RS exposure (RM ANOVA, ‘Treatment’ F_1,20_=4.62, *P* = 0.0441, *N* = 8/4). (**B**, **C**) Quantification of glucocorticoid receptor (GR) immunofluorescence intensity in the hippocampus of male (**B**, *N* = 7/8/6) and female (**C**, *N* = 9/9/5) mice, at one month after the MIA injection. (**D**) Representative images of GR staining in the hippocampus of male mice (scale bar 200μm). (**E**) Quantification of the total number of c-Fos positive neurons in the rostroventromedial medulla (RVM) of male mice at 2h after the MIA injection. *N* = 6/6/6 (**F**) Quantification (left) and representative images (right; scale bar = 100μm) of c-Fos immunopositive neurons in the locus coeruleus of male mice 2h after the MIA injection. *N* = 6/6/6.

**Figure S6.**
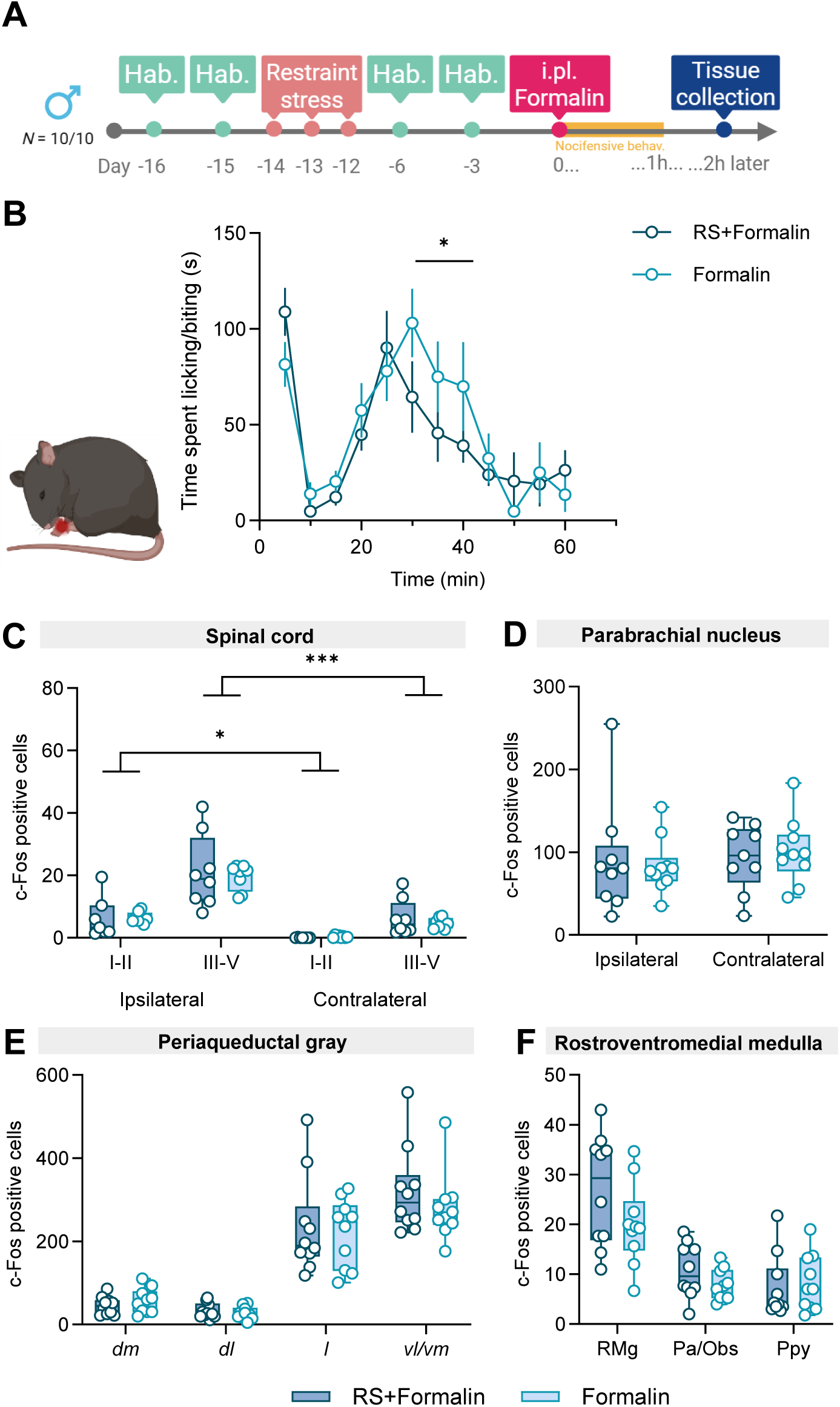
RS modifies the second phase of the formalin test but does not impact early c-Fos expression. (**A**) Timeline of experimental procedures (hab. – habituation, i.pl. – intraplantar). (**B**) Quantification of the time spent by male mice attending the hindpaw following the i.pl. injection of 5% formalin solution (2-way RM ANOVA, T30-45: F_1,17_=4.78, *P* = 0.0431). n.b., one of the mice *not* exposed to RS did not display the second phase of the nocifensive response, hence it was excluded from the analysis. Data presented as mean ± SEM. (**C-F**) Quantification of c-Fos immunopositive neurons at spinal (**C**: 2-way RM ANOVA: ‘Area’ F_3,39_=81.45, *P* < 0.0001; ‘Treatment’ F_1,14_=0.32, *P* = 0.5823; Šídák’s post-hoc) and supraspinal sites in male mice at two hours after the i.pl. injection of formalin. \**P* < 0.001, \*\*\*\**P* < 0.0001. *N* = 10/10 except for spinal cFos = 8/8)

**Figure S7.**
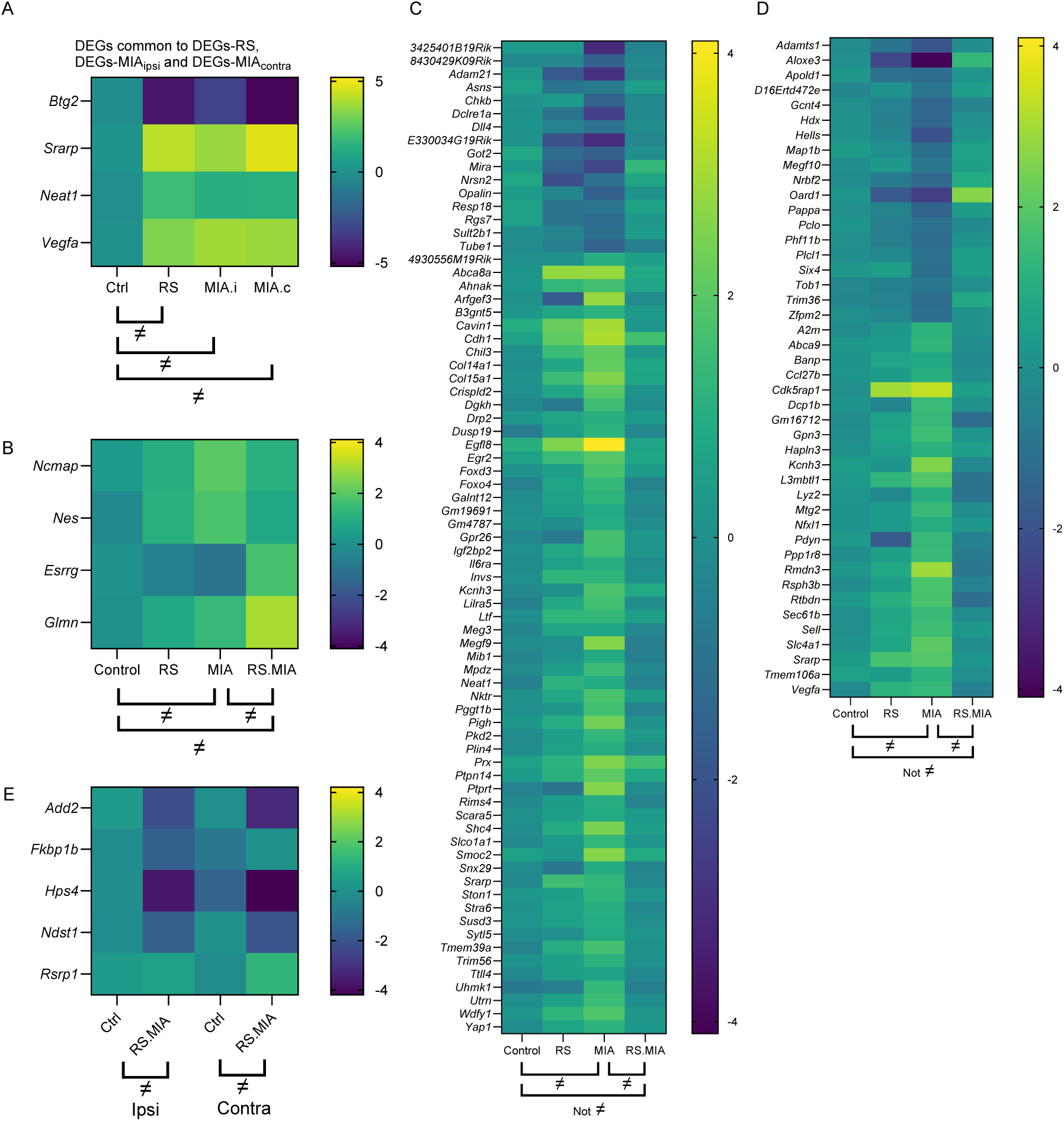
Shared and condition-specific transcriptomic responses to RS, MIA and RS+MIA. (**A**) Heatmap showing genes commonly differentially expressed across RS, MIA (ipsilateral), and MIA (contralateral) conditions. (**B)** Expression of genes differentially expressed across MIA, RS.MIA and MIA *vs* RS.MIA. Plots for *Ncmap* and *Nes* represent ipsilateral data while plots for *Esrrg* and *Glmn* are for the contralateral side, (**C, D**) MIA-induced gene changes mitigated by RS on the ipsilateral (**C, n=75**) and contralateral (**D, n=44**) spinal dorsal horn. (**E**) Heatmap of DEGs identified both ipsilaterally and contralaterally with a substantial impact of the combination RS + MIA. Statistical significance between conditions is indicated with “≠”; brackets denote comparisons.

**Table S1.** Description of statistical results for Figures 1 to 5.

| Fig. | Panel | N<br>m/f | Significa<br>nce Test | Factors /<br>Comparis<br>on | *F*, *t*, *R*<br>value | *P* value | Post-hoc test |
| --- | --- | --- | --- | --- | --- | --- | --- |
| 1 | B | 8/8/6<br>m | 2-way<br>RM<br>ANOVA<br>D-1 to 27 | Treatment | $F_{2,19}=8.18$ | $P = 0.0027$ | Šídák:<br>Control vs MIA: $P = 0.003$;<br>Control vs RS+MIA: $P = 0.018$. |
| | | | | Time | $F_{6,114}=2.64$ | $P = 0.0196$ | |
| | | | 2-way<br>RM<br>ANOVA<br>D-1 to 8 | Treatment | $F_{2,19}=5.49$ | $P = 0.013$ | Šídák:<br>Control vs MIA: $P = 0.011$<br>Control vs RS+MIA: $P = 0.303$. |
| | C | 8/8/6<br>m | 1-way<br>ANOVA | Treatment | $F_{2,19}=9.109$ | $P = 0.0017$ | Šídák (see in figure) |
| | D | 8/8/6<br>m | 2-way<br>RM<br>ANOVA<br>D-1 to 27 | Treatment | $F_{2,19}=11.67$ | $P = 0.005$ | Šídák:<br>Control vs MIA: $P < 0.001$<br>Control vs RS+MIA: $P = 0.052$. |
| | | | | Time | $F_{6,114}=1.87$ | $P = 0.0927$ | |
| | | | 2-way<br>RM<br>ANOVA<br>D-1 to 12 | Time | $F_{3,57}=3.42$ | $P = 0.023$ | |
| | | | 2-way<br>RM<br>ANOVA<br>D4 to 23 | Treatment | $F_{2,19}=14.9$ | $P < 0.001$ | Šídák:<br>Control vs MIA: $P < 0.001$;<br>MIA vs RS+MIA: $P = 0.039$. |
| | E | 8/8/6<br>m | 1-way<br>ANOVA | Treatment | $F_{2,19}=14.32$ | $P = 0.0002$ | Šídák (see in figure) |
| | F | 9/9/6<br>f | 2-way<br>RM<br>ANOVA<br>D-1 to 27 | Treatment | $F_{2,21}=8.95$ | $P = 0.0015$ | Šídák:<br>Control vs MIA $P = 0.079$;<br>Control vs RS+MIA: $P = 0.001$;<br>MIA vs RS+MIA: *ns*. |
| | | | | Time | $F_{6,126}=4.14$ | $P = 0.0008$ | |
| | | | 2-way<br>RM-ANOVA<br>D5 to 23 | Treatment | $F_{2,21}=10.64$ | $P < 0.001$ | Šídák:<br>RS+MIA vs Ct: $P < 0.001$;<br>MIA vs Control: $P = 0.037$. |
| | G | 9/9/6<br>f | 1-way<br>ANOVA | Treatment | $F_{2,21}=10.36$ | $P = 0.0007$ | Šídák (see in figure) |
| | H | 9/9/6<br>f | 2-way<br>RM<br>ANOVA<br>D-1 to 27 | Treatment | $F_{2,21}=6.83$ | $P = 0.0052$ | Šídák:<br>RS+MIA vs Control: $P = 0.008$<br>MIA vs Control: $P = 0.012$<br>MIA vs RS+MIA: *ns*. |
| | | | | Time | $F_{6,126}=1.353$ | $P = 0.2386$ | |
| | I | 9/9/6<br>f | 1-way<br>ANOVA | Treatment | $F_{2,21}=6.51$ | $P = 0.0063$ | Šídák (see in figure) |
| | J | 7 */8/6<br>m | 1-way<br>ANOVA | Treatment | $F_{2,18}=7.06$ | $P = 0.0055$ | Šídák (see in figure) |
| | K | 7 */8/6<br>m | 1-way<br>ANOVA | Treatment | $F_{2,18}=3.22$ | $P = 0.0636$ | Šídák:<br>RS+MIA vs MIA: $P = 0.314$;<br>RS+MIA vs Control: $P = 0.782$;<br>MIA vs Control: $P = 0.07$. |
| | L | 9/9/6<br>f | 1-way<br>ANOVA | Treatment | $F_{2,21}=11.44$ | $P = 0.0004$ | Šídák (see in figure) |
| | M | 9/9/6<br>f | 1-way<br>ANOVA | Treatment | $F_{2,21}=3.25$ | $P = 0.059$ | Šídák:<br>RS+MIA vs MIA $P = 0.8523$;<br>RS+MIA vs Control $P = 0.0596$; |
| | | | | | | | MIA vs Control $P = 0.2153$. |
| | O | 7*/8/6<br>m | 1-way<br>ANOVA | Treatment | $F_{2,19}=18.49$ | $P < 0.0001$ | Šídák (see in figure) |
| | P | 9/9/6<br>f | 1-way<br>ANOVA<br>Kruskal<br>Wallis | Treatment | $F_{2,21}=0.792$<br>Kruskal-Wallis<br>Statistics:<br>0.1700 | $P = 0.466$<br>$P = 0.9185$ | N/A |
| 2 | A | 8/7*/6<br>m | 1-way<br>ANOVA | Treatment | $F_{2,18}=5.15$ | $P = 0.017$ | Šídák (see in figure) |
| | B | 8/7*/6<br>m | 1-way<br>ANOVA<br>Kruskal<br>Wallis | Treatment | $F_{2,18}=5.97$<br>Kruskal<br>Wallis<br>statistics<br>11.88 | $P = 0.0103$<br>$P=0.006$ | Šídák(see in figure)<br>Kruskal Wallis:<br>RS+MIA vs MIA: $P = 0.0039$;<br>MIA vs Control: $P = 0.0223$ |
|  | C | 8/7*/6<br>m | 1-way<br>ANOVA | Treatment |  | ns |  |
|  | D | 8/7*/6<br>m | 1-way<br>ANOVA | Treatment |  | ns |  |
| | F | 8*/9/6<br>f | 1-way<br>ANOVA | Treatment | $F_{2,21}=3.09$ | $P = 0.067$ | N/A |
| | G | 8*/9/6*<br>f | 1-way<br>ANOVA | Treatment | $F_{2,20}=2.99$ | $P = 0.0731$ | N/A |
| | H | 8*/9/6<br>f | 1-way<br>ANOVA | Treatment | $F_{2,20}=3.8$ | $P = 0.0399$ | Šídák (see in figure) |
| | I | 8*/9/6<br>f | 1-way<br>ANOVA | Treatment | $F_{2,20}=4.36$ | $P = 0.0269$ | Šídák (see in figure) |
| | J | 8/7*/6<br>m | Simple<br>linear<br>regression | | J1: $R^2=0.2143$<br>$F_{1,19}=5.18$<br>J2: $R^2=0.1765$<br>$F_{1,19}=4.07$ | J1: $P = 0.0346$<br>J2: $P = 0.058$ | N/A |
| | K | 8*/9/6<br>f | Simple<br>linear<br>regression | | K1: $R^2=0.1828$<br>$F_{1,21}=4.7$<br>K2: $R^2=0.1936$<br>$F_{1,21}=5.04$ | K1: $P = 0.0418$<br>K2: $P = 0.0357$ | N/A |
| | L | 8/8/6 | 2-way<br>RM<br>ANOVA | Time | $F_{1,19}=90.90$ | $P < 0.0001$ | Šídák (see in figure) |
| | M | 9/9/6 | 2-way<br>RM<br>ANOVA | Time | $F_{1,21}=44.52$ | $P < 0.0001$ | Šídák (see in figure) |
| 3 | A | 8/6/6<sup>\$</sup> | 2-way<br>RM<br>ANOVA | Treatment | $F_{2,17}=3.54$ | $P = 0.050$ | Šídák (see in figure) |
|  | C | 9/9/6 | 2-way<br>RM<br>ANOVA | Treatment |  | ns |  |
| | D | 6/8/5<sup>#</sup> | 1-way<br>ANOVA | Treatment | $F_{2,16}=5.21$ | $P = 0.0181$ | Šídák (see in figure) |
|  | F | 8/9/5<sup>#</sup> | 1-way<br>ANOVA | Treatment |  | ns |  |
| | G | 7/8/6<sup>\$</sup> | 2-way RM ANOVA | Treatment | | *ns* | |
|  | I | 8/9/5<sup>#</sup> | 2-way RM ANOVA | Treatment |  | *ns* |  |
| | J | 7/8/6<sup>\$</sup> | 2-way RM ANOVA | Treatment | | *ns* | |
|  | K | 8/9/5<sup>#</sup> | 2-way RM ANOVA | Treatment |  | *ns* |  |
| 4 | A | 6/4<sup>\$</sup>/4<sup>\$</sup>m | 3-way ANOVA | Treatment | $F_{2,44}=6.875$ | $P = 0.003$ | Šídák<br>MIA vs RS + MIA: $P = 0.002$. |
| | | | | Side | $F_{1,44}=20.642$ | $P < 0.001$ | |
| | | | | Laminae | $F_{1,44}=178.792$ | $P < 0.001$ | |
| | | | | Treat x Laminae | $F_{2,44}=8.824$ | $P < 0.001$ | |
| | | | Post-hoc 2-way ANOVA Laminae I, II | Treatment | $F_{2,22}=5.124$ | $P = 0.015$ | Šídák<br>Control vs MIA: $P = 0.026$;<br>Control vs RS+MIA: $P = 0.035$ |
| | | | | Side | $F_{1,22}=35.059$ | $P < 0.001$ | |
| | | | | Treat x Side | $F_{2,22}=3.55$ | $P = 0.046$ | |
| | | | Post-hoc 2-way ANOVA Laminae III-V | Treatment | $F_{2,22}=8.052$ | $P = 0.002$ | Šídák<br>Control vs RS+MIA: $P = 0.020$;<br>MIA vs RS+MIA: $P = 0.004$. |
| | | | | Side | $F_{1,22}=9.578$ | $P = 0.005$ | |
| | | | Post-hoc 1-way ANOVA Laminae I,II ipsi | Treatment | Šídák | $P = 0.033$ | Šídák<br>RS+MIA vs Control: $P = 0.041$. |
| | | | Post-hoc 1-way ANOVA Laminae III-V ipsi | Treatment | Šídák | $P = 0.021$ | Šídák<br>RS+MIA vs MIA: $P = 0.022$. |
| | B | 6/6/6m | 2-way RM ANOVA | Treatment | $F_{2,15}=4.6$ | $P = 0.0276$ | Šídák (see in figure) |
| | | | | Area | $F_{1,15}=34.73$ | $P < 0.0001$ | Šídák (see in figure) |
| | C | 6/6/6m | 2-way RM ANOVA | Treatment | $F_{2,1}=4.54$ | $P = 0.0287$ | Šídák (see in figure) |
| | | | | Side | $F_{1,15}=3.18$ | $P = 0.095$ | N/A |
| | D | 6/6/6m | 1-way ANOVA | Treatment | $F_{2,15}=4.52$ | $P = 0.0291$ | Šídák (see in figure) |

\* outlier identified using the Tukey's IQR test in at least 2 outcome measures

**Table S2.**
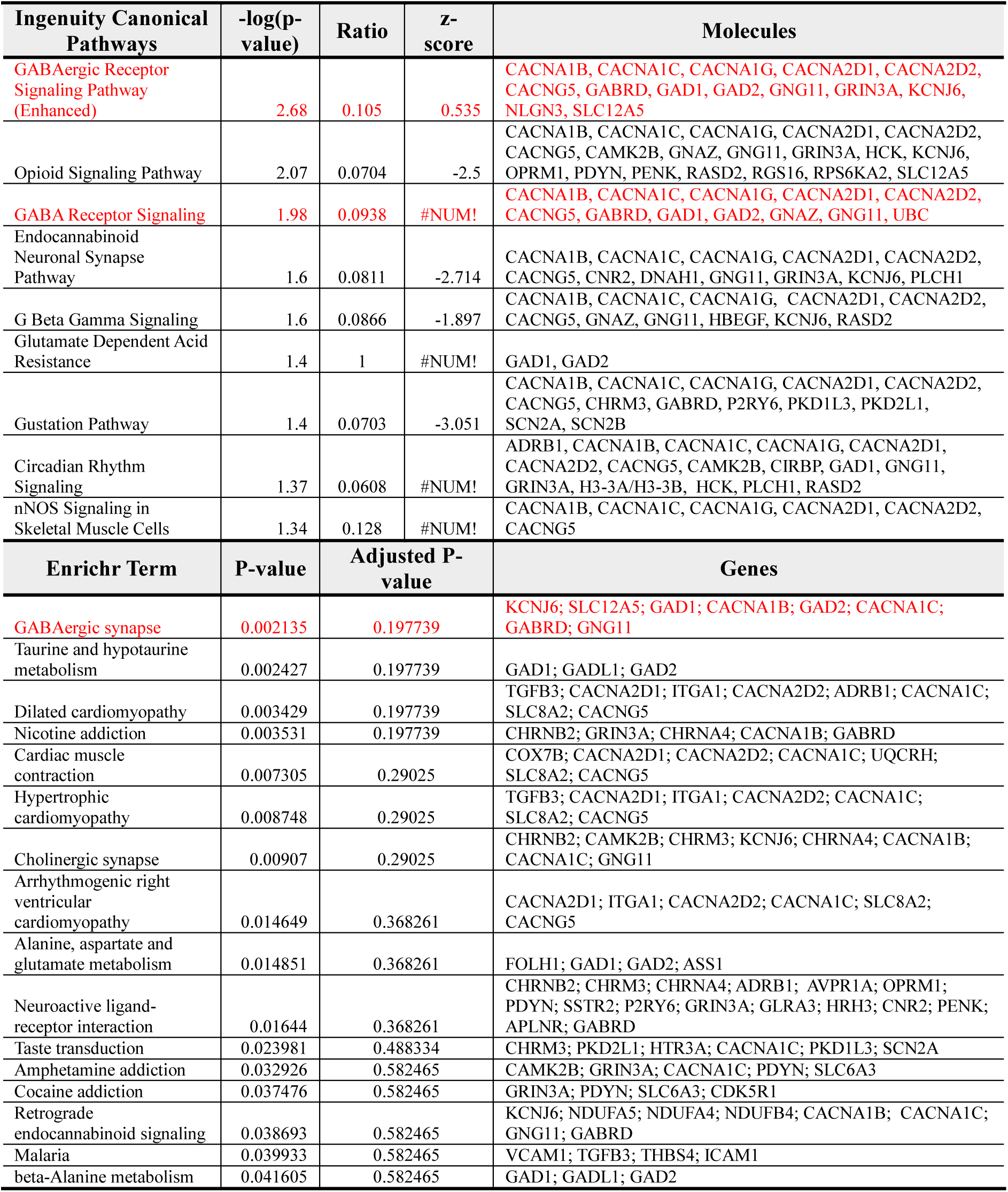
Pathway analysis for DEGs identified by the RS *vs* Control contrast. IPA analysis threshold for reporting: *P* < 0.05 or –log(p-value) = 1.30. Enrichr threshold for reporting: *P* < 0.05.

**Table S3.**
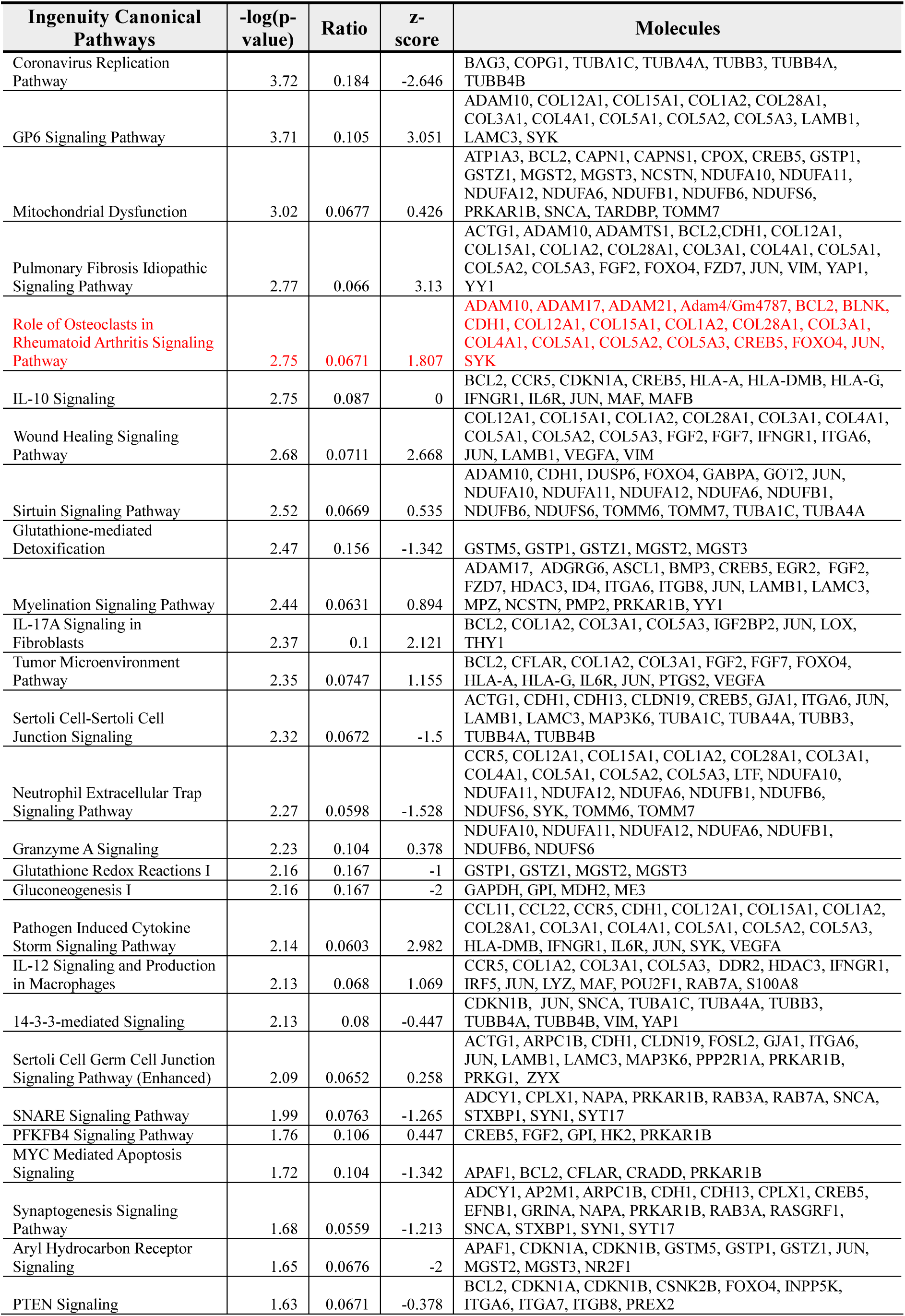
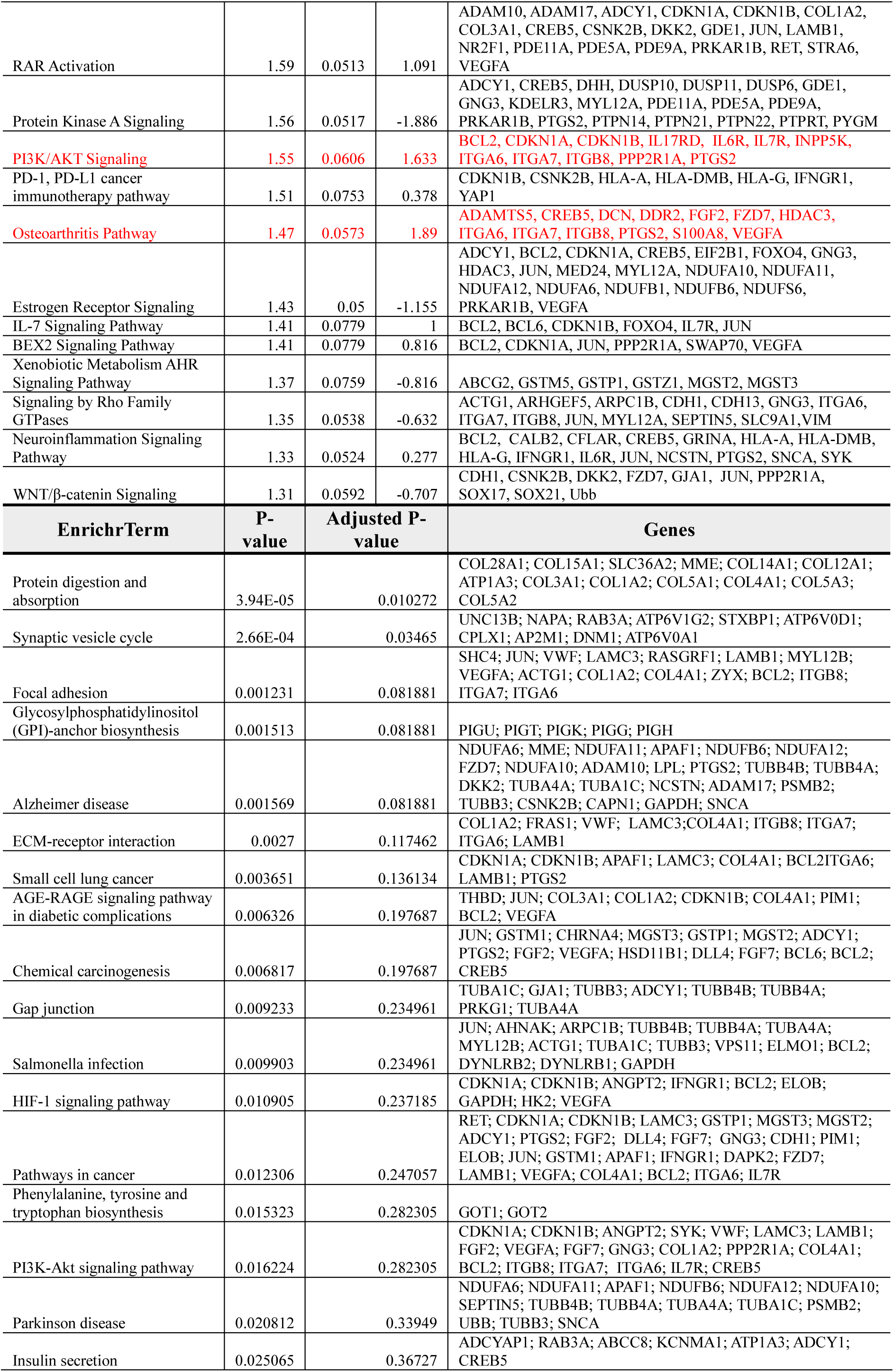
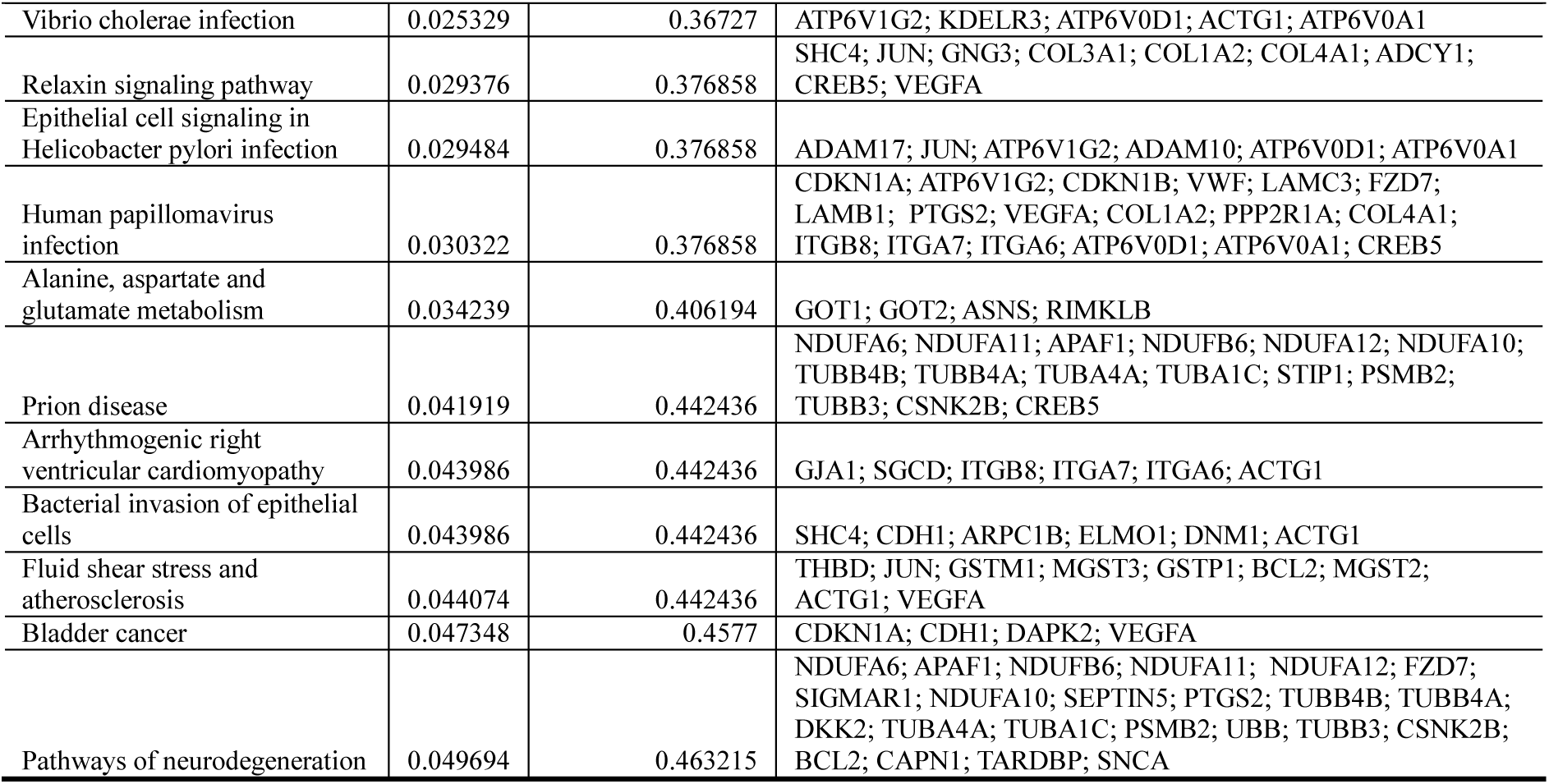
Pathway analysis for DEGs identified by the MIA vs Control ipsi contrast. IPA analysis threshold for reporting: *P* < 0.05 or –log(p-value) = 1.30. Enrichr threshold for reporting: *P* < 0.05.

**Table S4.** Pathway analysis for DEGs identified by the MIA vs Control contra contrast. IPA analysis threshold for reporting: P<0.05 or –log(p-value) = 1.30. Enrichr threshold for reporting: *P* < 0.05.

| Ingenuity Canonical Pathways | $-\log(p\text{-value})$ | Ratio | z-score | Molecules |
| --- | --- | --- | --- | --- |
| Pulmonary Healing Signaling Pathway | 2.76 | 0.0359 | 0.378 | FZD5,HCK,MMP19,NOTCH2,PRKCH,VEGFA,WNT5B |
| WNT/Ca<sup>+</sup> pathway | 2.57 | 0.0606 | -1 | CREB5,FZD5,PLCL1,WNT5B |
| Regulation of the Epithelial Mesenchymal Transition in Development Pathway | 2.17 | 0.0471 | 0 | FZD5,GLIS1,TWIST1,WNT5B |
| Pulmonary Fibrosis Idiopathic Signaling Pathway | 2.14 | 0.0252 | -1.414 | ADAMTS1,CCN2,COL19A1,FZD5,MMP19,NOTCH2,PLAU,WNT5B |
| Factors Promoting Cardiogenesis in Vertebrates | 2 | 0.0338 | -0.447 | CREB5,FZD5,PLCL1,PRKCH,WNT5B |
| Osteoarthritis Pathway | 1.81 | 0.0264 | 1 | CASP6,CREB5,FZD5,GLIS1,S100A9,VEGFA |
| Role of Macrophages, Fibroblasts and Endothelial Cells in Rheumatoid Arthritis | 1.67 | 0.0223 | #NUM! | CREB5,FZD5,PLCL1,PRKCH,SFRP4,VEGFA,WNT5B |
| Basal Cell Carcinoma Signaling | 1.64 | 0.0435 | #NUM! | FZD5,GLIS1,WNT5B |
| Axonal Guidance Signaling | 1.51 | 0.0183 | #NUM! | ADAMTS1,EFNA4,FZD5,GLIS1,PAPPA,PLCL1,PRKCH,VEGFA,WNT5B |
| Chronic Myeloid Leukemia Signaling | 1.47 | 0.022 | 0 | FZD5,GLIS1,HCK,NOTCH2,PLCL1,WNT5B |
| Endocannabinoid Cancer Inhibition Pathway | 1.42 | 0.028 | 1 | CASP6,CREB5,TWIST1,VEGFA |
| Coagulation System | 1.42 | 0.0571 | #NUM! | A2M,PLAU |
| Inhibition of Matrix Metalloproteases | 1.38 | 0.0541 | #NUM! | A2M,MMP19 |
| Galactose Degradation I (Leloir Pathway) | 1.36 | 0.2 | #NUM! | GALM |
| EnrichrTerm | P-value | Adjusted P-value | Genes |  |
| RNA degradation | 8.52E-04 | 0.1073 | BTG2;PABPC5;TOB2;TOB1;DCP1B |  |
| Transcriptional misregulation in cancer | 0.03389 | 0.8666 | SIX4;PLAU;FLI1;DUSP6;MLF1 |  |
| SNARE interactions in vesicular transport | 0.03762 | 0.8666 | BNIP1;STX11 |  |
| Proteoglycans in cancer | 0.04294 | 0.8666 | WNT5B;FZD5;PLAU;TWIST1;VEGFA |  |
| Insulin secretion | 0.04621 | 0.8666 | PCLO;KCNN2;CREB5 |  |
| GABAergic synapse | 0.05027 | 0.8666 | PLCL1;GABRG3;GABRD |  |

**Table S5.** Pathway analysis for DEGs identified in both Ctrl *vs* RS+MIA and MIA *vs* RS+MIA, on the ipsilateral side. Enrichr threshold for reporting: *P* < 0.05.

| Term | P-value | Adjusted P-value | Genes |
| --- | --- | --- | --- |
| Transcriptional misregulation in cancer | 0.023655 | 0.497236 | CCNT2;EYA1;KLF3 |
| Ubiquinone and other terpenoid-quinone biosynthesis | 0.03465 | 0.497236 | GGCX |
| Longevity regulating pathway | 0.042318 | 0.497236 | SESN3;CREB3L2 |
| Glucagon signaling pathway | 0.046121 | 0.497236 | CREB3L2;SIK2 |

**Table S6.**
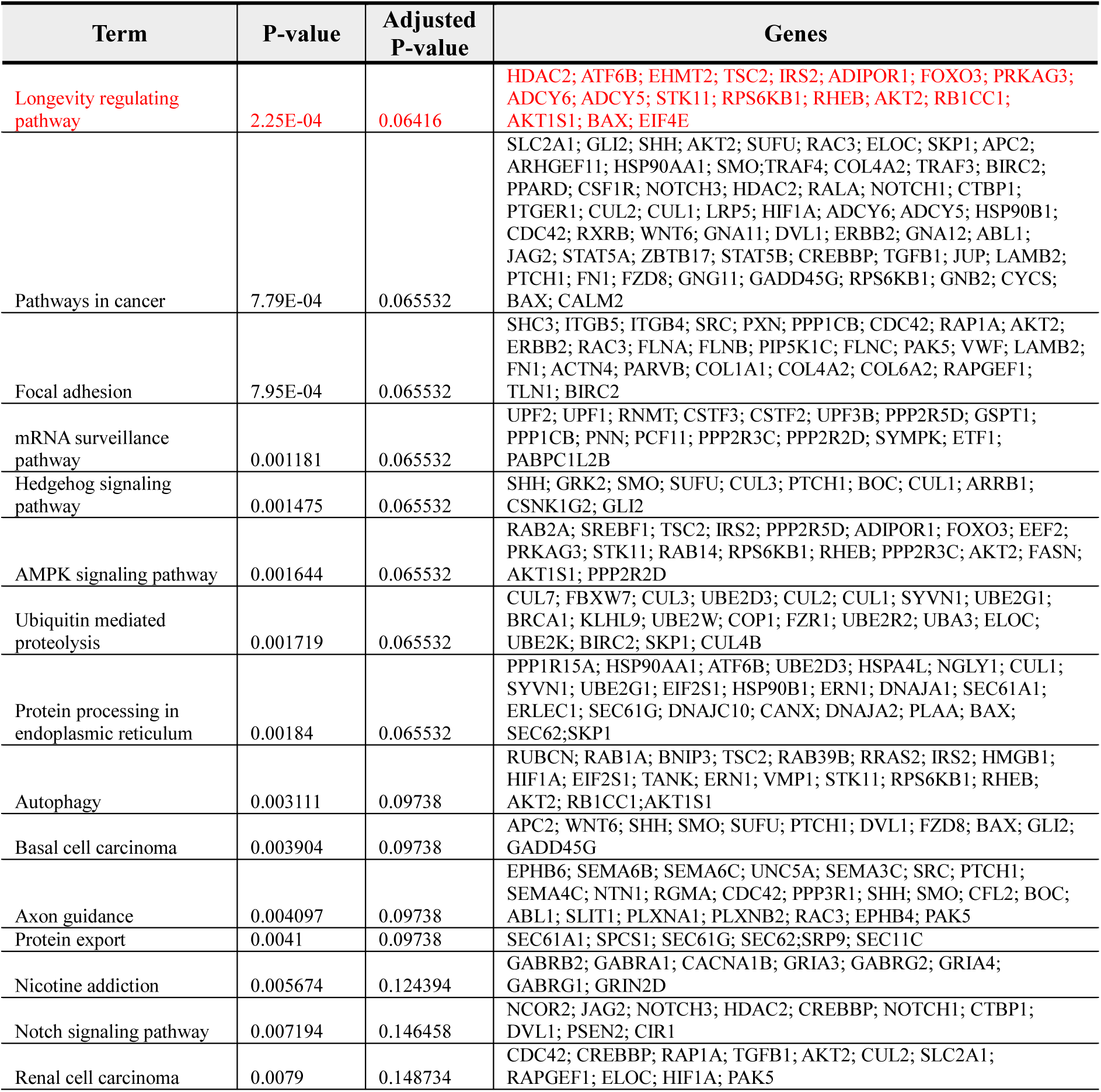
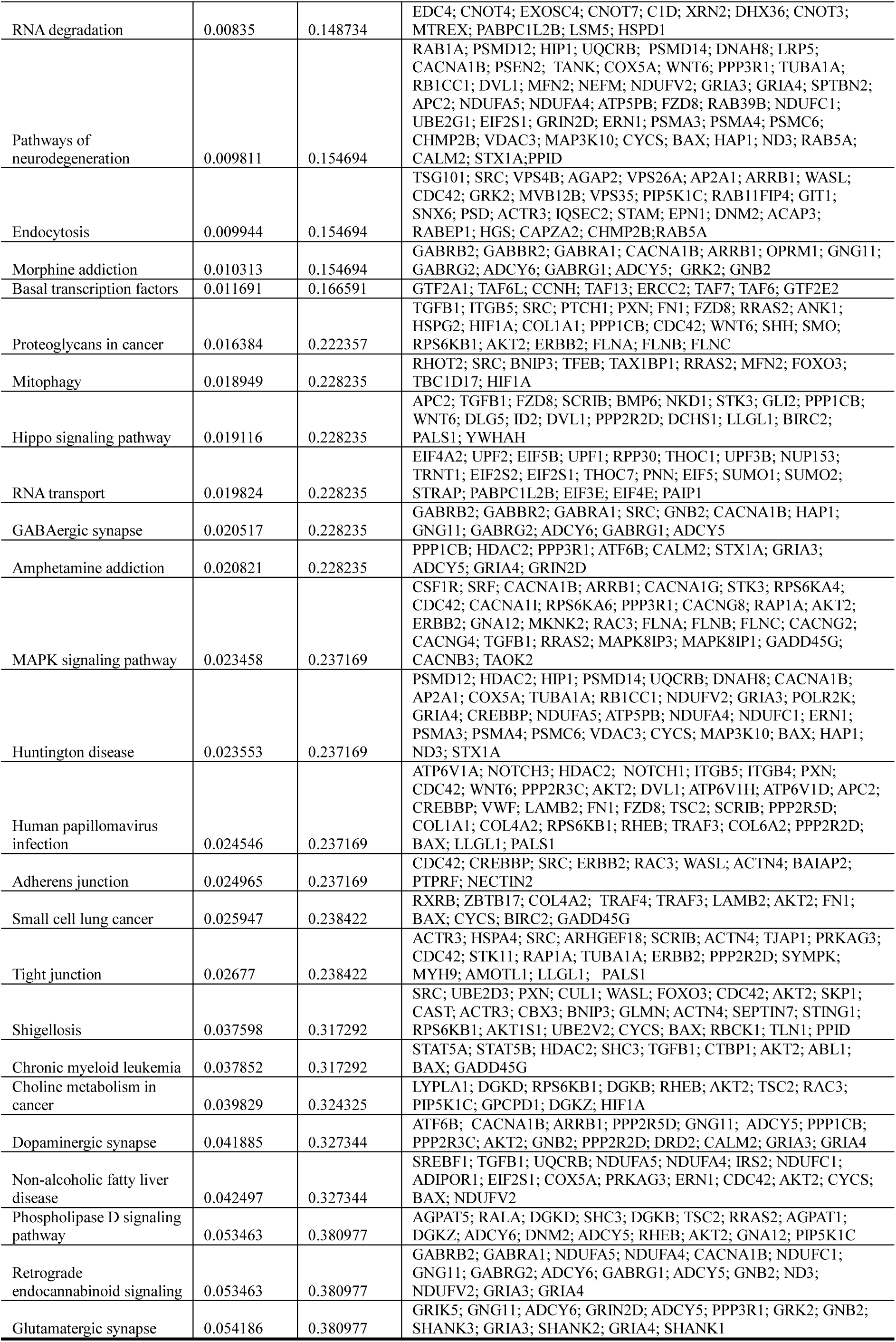
Pathway analysis for DEGs identified in both Ctrl *vs* RS+MIA and MIA *vs* RS+MIA, on the contralateral side. Enrichr threshold for reporting: *P* < 0.055.

